# Mechanical stress in pancreatic cancer: Signaling pathway adaptation activates cytoskeletal remodeling and enhances cell migration

**DOI:** 10.1101/2021.06.11.448065

**Authors:** Maria Kalli, Ruxuan Li, Gordon B. Mills, Triantafyllos Stylianopoulos, Ioannis K. Zervantonakis

**Author notes:** These authors contributed equally to this work (Maria Kalli, Ruxuan Li). Corresponding authors, **Address correspondence to:** University of Pittsburgh, Department of Bioengineering, 300 Technology Dr., Room 309, Pittsburgh, PA, 15219, United States., University of Cyprus, Cancer Biophysics Laboratory, Mechanical and Manufacturing Engineering, 75 Kallipoleos St., Nicosia, 1678, Cyprus.

## Abstract

New treatments for patients with advanced or metastatic pancreatic cancers are urgently needed due to their resistance to all current therapies. Current studies focus on alternative treatment approaches that target or normalize the abnormal microenvironment of pancreatic tumors, which among others, is responsible for elevated mechanical stress in the tumor interior. Nevertheless, the underlying mechanisms by which mechanical stress regulates pancreatic cancer metastatic potential remain elusive. Herein, we used a large-scale proteomic assay to profile mechanical stress-induced signaling cascades that drive the motility of pancreatic cancer cells. Proteomic analysis, together with selective protein inhibition and siRNA treatments, revealed that mechanical stress enhances cell migration through activation of the p38 MAPK/HSP27 and JNK/c-Jun signaling axes, and activation of the actin cytoskeleton remodelers: Rac1, cdc42, and Myosin II. Our results highlight targeting aberrant signaling in cancer cells that are adapted to the mechanical tumor microenvironment as a novel approach to effectively limit pancreatic cancer cell migration.

## Introduction

Pancreatic ductal adenocarcinoma (PDAC) is responsible for over 90% of pancreatic cancer cases and is the seventh leading cause of cancer-related death in the United States^1^. PDAC is characterized by extremely poor prognosis with a 5-year survival rate of less than 8%^1^. New treatments for pancreatic cancer are urgently needed as it is highly resistant to all current chemotherapy and radiotherapy approaches. Complete surgical resection is currently the only curative treatment, however, patients with advanced or metastatic pancreatic cancer are most often ineligible for surgical removal, and even following resection, the tumor usually relapses within a year after surgery^1^.

Current studies focus on normalizing the microenvironment of PDACs^2, 3^. The PDAC microenvironment is characterized by a dense tumor-associated stroma composed of extracellular proteins, such as collagen I and hyaluronan that are remodeled to create a stiff Extracellular Matrix (ECM), a condition known as desmoplasia^4–7^. Losartin, through its TGFβ inhibitory activity, has been tested in the clinic as an approach to target the dense stroma^8^. Increased matrix stiffness also serves as a diagnostic marker and is associated with poor prognosis. It can activate focal adhesion proteins, such as focal adhesion kinase (FAK) and paxillin, and actin cytoskeleton remodelers, such as RAC, RHO GTPase/ Rho-associated kinase (ROCK) and RAS GTPases, to trigger signaling cascades that induce cell motility, migration and invasion^9–11^. Moreover, tumor growth within a physically-restricted environment also leads to the development of mechanical compressive forces in the tumor interior, resulting in intra-tumoral compressive stresses that can reach up to 75mmHg (10kPa)^12–15^. This type of mechanical stress has been shown to activate signaling pathways that promote tumorigenesis and invasiveness ^16–18^. However, the molecular mechanisms that underly the effects of mechanical stress on metastatic potential remain elusive.

We hypothesized that mechanical stress activates signaling pathways shown to be consistently activated in pancreatic tumors. These pathways include the PI3K/Akt and Ras/MAPK signaling cascades that are involved in the regulation of cell survival and motility ^12^. The increased Ras/MAPK signaling is likely due to K-Ras being the most frequently mutated gene in invasive pancreatic tumors with a rate of 95%^3^. K-Ras can activate the PI3K/Akt pathway, but its main target is the MAPK signaling cascade, that includes the Jun N-terminal kinase (JNK) and p38 MAPK. These kinases are normally activated by environmental and genotoxic stresses and play key roles in the regulation of cell proliferation, survival and migration. Although it less clear, they could also contribute to mechanical stress-induced signal transduction^19^. In our previous studies, we showed that mechanical stress promotes GDF15-induced cancer cell migration through activation of PI3K/Akt and MEK1/Erk1 signaling cascades in pancreatic and brain cancer cell lines, respectively ^20, 21^. Based on these findings, we used a large-scale proteomic assay to identify mechanical stress-induced signaling cascades that could contribute to motility of pancreatic cancer cells. Our results elucidate mechanical stress-induced signaling mechanisms and identify therapeutic targets to limit pancreatic cancer cell migration.

## Results

### Proteomic profiling reveals heterogeneous responses across multiple pathways and uncovers activation of p38 MAPK/HSP27 and JNK/c-Jun as a mechanical stress-induced signaling pathway in pancreatic cancer cells

To profile the mechanical stress-induced signaling pathway adaptation in pancreatic cancer cells, we analysed total and phosphorylated protein levels following application of a compressive force on the cells. Specifically, 425 proteins including phospho-proteins were assessed using Reverse Phase Protein Arrays (RPPA), from which around 95 resulted in a statistically significant change in the compressed compared to the uncompressed cells. Proteins with the largest fold-change (20%) increases (red) or decreases (blue) between control and compressed cells were visualized in a heatmap (**Figure 1A, Supplementary Figure 1, Appendix I**). We found that mechanical stress activated a number of regulators that mediate cellular adaptation to multiple stress signals, including heat shock stress, endoplasmic reticulum stress, inflammatory stress and oxidative stress. Specifically, the cellular stress response mediator, Heat Shock Protein 27 (HSP27_pS82) and its upstream activator p38 Mitogen Activated Protein Kinase (p38 MAPK_pT180_Y182), exhibited a strong activation in compressed compared to control cells, that was also validated through Western Blotting in both MIA PaCa-2 and PANC-1 cells (**Figure 1 -B, -C, and -E)**. In line with p38 MAPK activation, we found that the downstream target of the stress response protein c-Jun N-terminal Kinase (JNK), c-Jun (pS73), was also activated in compressed cells. Subsequently both c-Jun and JNK activation were validated through Western Blotting (**Figure 1D-E, Supplementary Figure 2**). Additional markers of cellular stress, including the LC3A-B, a marker of autophagosomes, and the Endoplasmic Reticulum (ER) stress sensor chaperone BiP-GRP78 (also known as Heat Shock Protein 70), were elevated in response to compression (**Figure 1A**). Among the top 10 upregulated proteins, we observed a strong upregulation of the Plasminogen Activator Inhibitor-1 (PAI-1) that is implicated in actin cytoskeleton organization and cell contraction. The effects of PAI-1 are mediated by downstream activation of Myosin II, that interacts with actin filaments to form actomyosin, allowing cell contraction and thus, cell motility and invasion^20, 22–26^. Indeed, Myosin II was also activated in compressed cells (Myosin II_pS1943) as shown in **Figure 1A**. The increase in expression and phosphorylation of these mediators could contribute to the enhanced migration of cells under mechanical compression^20^. On the other hand, mechanical stress downregulated multiple targets of the mammalian Target of Rapamycin (mTOR) pathway, including the p70-S6K_pT389, Rictor_pT1135, 4E-BP1_pS65 and mTOR_pS2448, which promote protein synthesis and cell growth and proliferation (**Figure 1A**). To evaluate how mechanical stress alters signaling pathways, we computed pathway scores for 9 major pathways ^27^. We found that multiple protein members of the cell cycle pathway (p27_pT157, Cyclin, E1,- B1 and -D1) and the TSC/mTOR pathway were reduced. In contrast, protein members of the PI3K/Akt (Akt_pS473, GSK3β_pS9, p27_pT198) and the Ras/MAPK (MEK1_pS217, MAPK_pT202_Y204, p90RSK_pT573, JNK_pT183_Y185, c-Jun_pS73) signaling pathways were elevated (**Figure 1F**, **Supplementary Figure 3)**. Collectively, our results show that mechanical stress activates cellular stress response mediators and actin cytoskeleton regulators that may trigger cell motility and impair cell cycle progression and proliferation.

**Figure 1.**
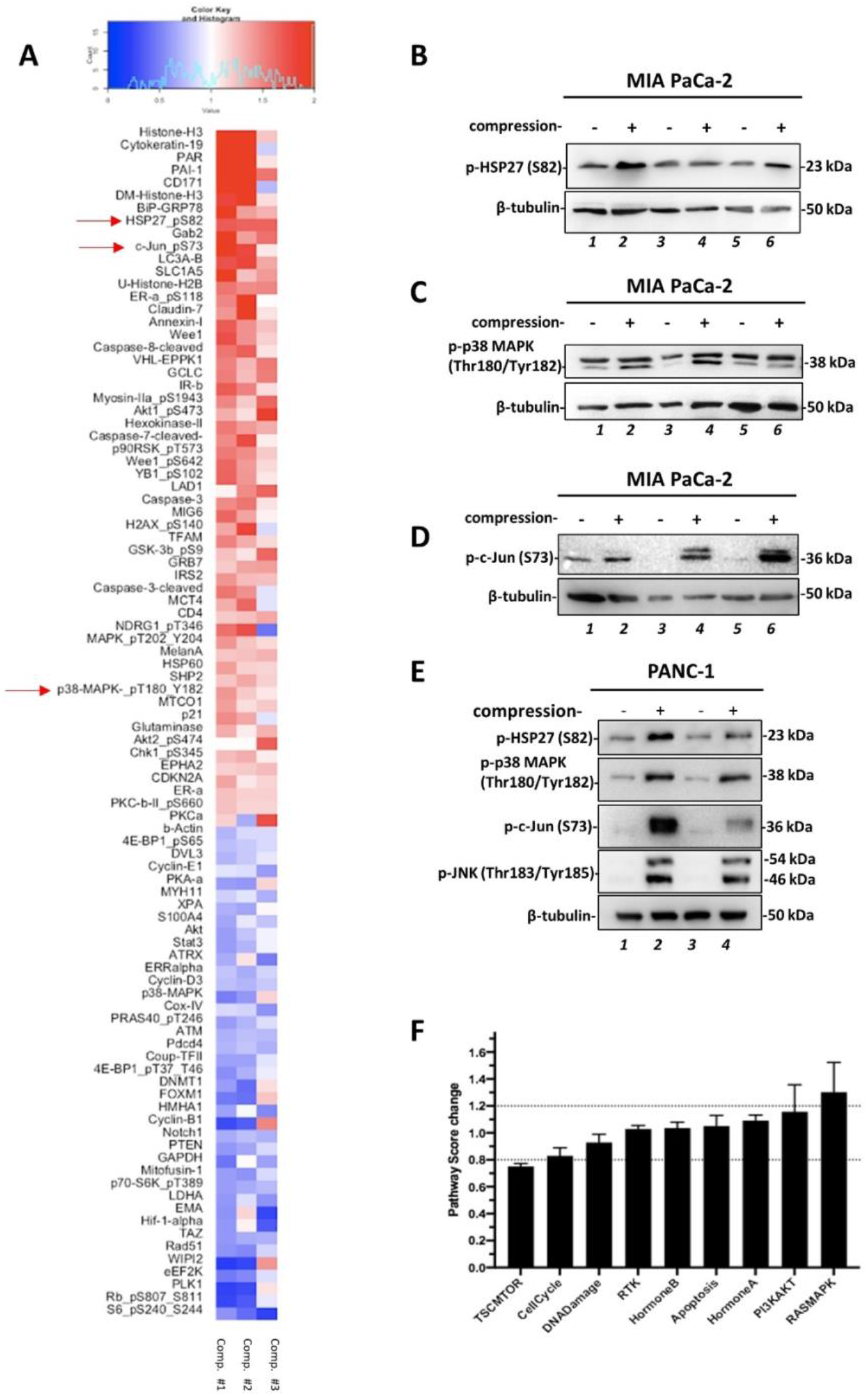
A Reverse Phase Protein Array was performed to reveal the mechanical stress induced mechanism in pancreatic cancer cell lines. **A,** Heatmap showing proteins exhibiting the largest fold increases (red) or decreases (blue) in compressed MIA PaCa-2 pancreatic cancer cells relative to the control cells. Proteins were analyzed by RPPA and paired t-test was performed to identify proteins that are differently expressed between compressed and uncompressed cells. Values in the heatmap represent the log2-ratio (compressed/uncompressed) of expression level for each protein in each condition (n=3; 3 biological replicates). Differences were considered significant with p < 0.05. **B-D**, Western Blotting was performed using the same lysates (MIA PaCa-2) that were analysed by RPPA to validate the activation of phospho-HSP27 (S82) **(B)**, phospho-p38 MAPK (Thr180/Tyr182) **(C)** and phospho-c-Jun (S73) **(D)**. **E**, Validation of phospho-JNK, in PANC-1 pancreatic cancer cells from 2 biological replicates. **F**, Pathway score analysis was calculated as the average sum of expression level of all members in each pathway, and then normalized to the uncompressed expression level. The protein members of the pathways are shown in Supplementary Figure 2.

### Mechanical stress induces actin cytoskeleton remodelling and contractility of pancreatic cancer cells promoting their migratory ability

Changes in the actin cytoskeleton organization in compressed MIA PaCa-2 and PANC-1 pancreatic cancer cell lines were evaluated using phalloidin staining. We observed increased stress fibres, filopodia and lamellipodia formation in both compressed MIA PaCa-2 and PANC-1 (**Figure 2A-B**) compared to uncompressed cells. In line with this, and based on Myosin II (Myosin IIa_pS1943) activation shown in RPPA **(Figure 1A)**, we also performed phospho-Myosin II staining and found increased phospho-Myosin II levels in both compressed MIA PaCa-2 and PANC-1 cells. Phospho-Myosin II was also co-localized with actin filaments in compressed cells (**Figure 2C-D**) suggesting increased actomyosin contractility. To evaluate whether these changes in cytoskeleton facilitate cell motility under mechanical stress, we performed a scratch wound assay. We found that compressed PANC-1 cells exhibited increased migratory ability as compared to the uncompressed cells (**Figure 2E-F**), similar to the effect that was previously observed in MIA PaCa-2 cells ^20^. Based on the findings of elongated cell shape and increased migration, we analysed the expression of epithelial to mesenchymal transition (EMT) genes *Snail, Slug* and *Twist* with qPCR. These genes were consistently upregulated in both cell lines, further verifying the enhanced motility of compressed cells (**Figure 2G**). Finally, we also examined the activation of Rac1 and cdc42 GTPases that regulate actin cytoskeleton organization for the formation of cell protrusions and actomyosin contractility by inducing Myosin II activation^23, 37, 38^. Indeed, G-LISA showed that the Rac-1 small GTPase is activated as early as 30 minutes after exposure of cells to compression, while the cdc42 GTPase is activated at 16 hours (**Figure 2H**), suggesting a possible mechanical stress-induced activation of the Rac1/cdc42/Myosin II axis. At the same time, the cell proliferation indicators AlamarBlue assay and Ki67 staining did not show significant changes in compressed compared to control MIA PaCa-2 and PANC-1 cells for at least 16 hours post-compression (**Supplementary Figure 4**). Although most of the protein targets of the cell cycle showed a decrease in their phosphorylation levels in cells compressed for 16 hours, quantifiable changes in cell proliferation were only observed 48 hours post-compression (**Supplementary Figure 4).** Therefore, our results suggest that mechanical stress can induce changes in actin cytoskeleton through a *Rac1*/cdc42/myosin II axis and gene expression alterations to trigger cell migration.

**Figure 2.**
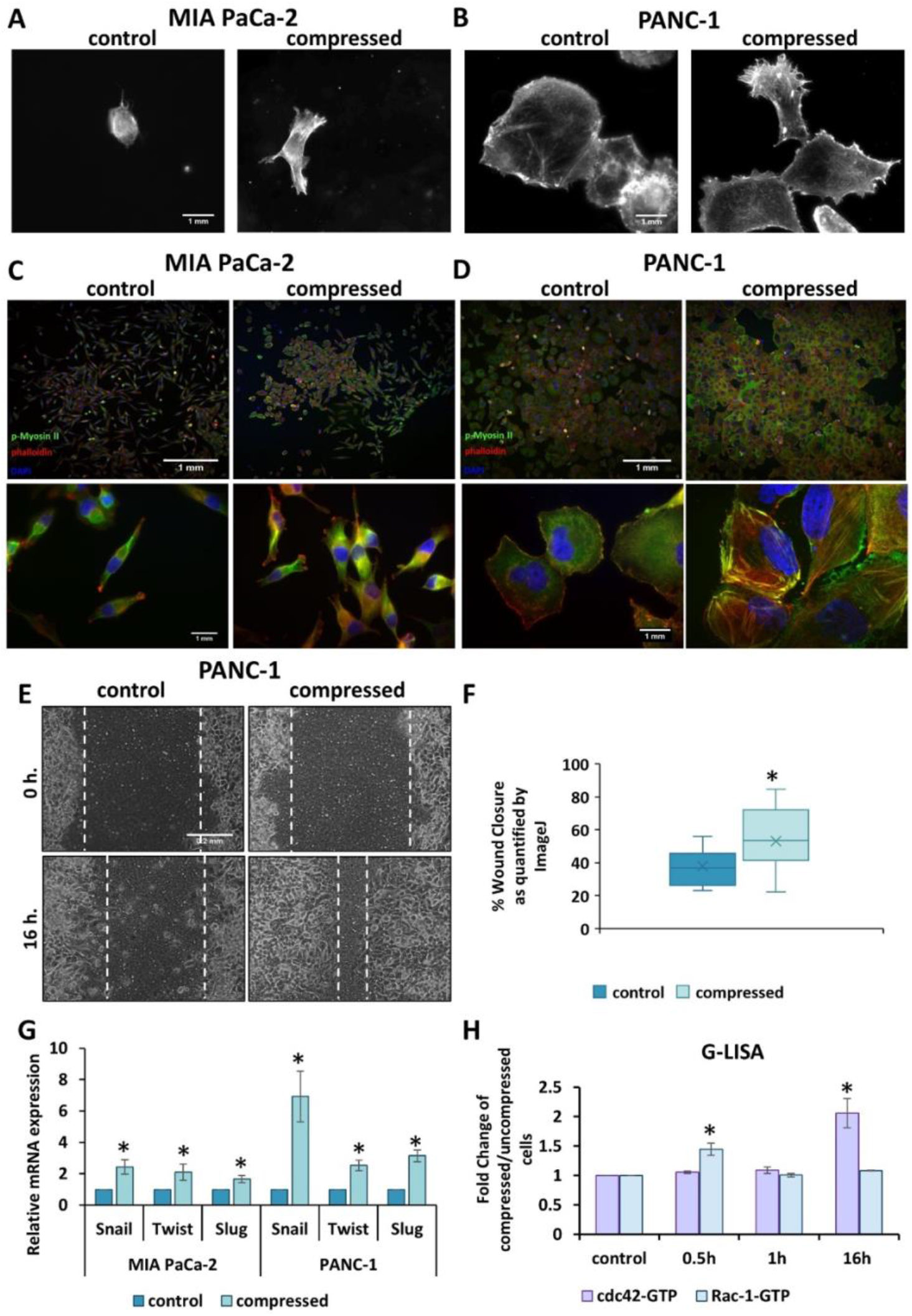
Mechanical stress induces cytoskeletal changes and cell contraction in pancreatic cancer cells promoting their migratory ability. **A-B,** Phalloidin staining was performed to monitor cell shape, actin cytoskeleton organization of control (0 mmHg) and compressed (4 mmHg) MIA PaCa-2 (A) and PANC-1 (B) cancer cells. Scale bar 1 mm. **C-D**, Staining for phospho-Myosin II (Ser1943) was performed to show actomyosin contractility in control and compressed MIA PaCa-2 (C) and PANC-1 (D) cells. Scale bar 1mm. **E,** PANC-1 cells were compressed by 4.0 mmHg in low-serum medium and then subjected to a scratch wound healing assay for 16 hours. Control cells (0 mmHg) were compressed by an agarose cushion only. Scale bar: 0.2 mm. **F,** Graph showing the average percentage of wound closure ±SE as quantified using ImageJ software. Statistically significant difference in wound closure of compressed PANC-1 cells compared to control cells is indicated with an asterisk (*) (n≥10; 3 biological replicates; p<0.05 in student’s t test). **G**, Relative mRNA expression of EMT markers *Snail, Twist, Slug* as quantified by qPCR in MIA PaCa-2 and PANC-1 cells. Each bar indicates the mean fold change ±SE of three independent experiments (n=9). Asterisk (*) indicates a statistically significant difference (p<0.05 in student’s t test). **H**, G-LISA was performed to analyze activation of cdc42- and Rac-1– small GTPases in compressed (4.0 mmHg) cells at different time points. Assay was performed in triplicates and graphs represent the average fold change ±SE of each protein in compressed relative to uncompressed (control) cells. Asterisk (*) indicates a statistically significant difference (p<0.05 in student’s t test).

### The p38 MAPK/HSP27 and JNK/c-Jun pathways are necessary for proliferation and migration of pancreatic cancer cells under mechanical stress

To evaluate whether the activation of the stress response pathway, p38 MAPK/HSP27, and JNK/c-Jun played a role in the increased migratory potential of cells under mechanical stress, we treated MIA PaCa-2 and PANC-1 cells with p38 MAPK (SB202190) or JNK (SP600125) inhibitors, and performed a scratch wound assay. We observed a significant decrease in the migration of MIA PaCa-2 and PANC-1 cancer cells when treated with either inhibitor under compression (**Figure 3A-C**). Cell proliferation was also effectively reduced by either inhibitor, as revealed by Ki67 staining (quantification in **Figure 3D-E, Supplementary Figure 5D**). Western blotting for phospho-HSP27 and phospho-c-Jun, that are downstream of p38 MAPK and JNK respectively, was also performed to confirm effective pathway inhibition for each inhibitor **(Figure 3F, Supplementary Figure 5A**). To further support our conclusion, we evaluated EMT markers using qPCR and observed that the expression of *Slug* and *Snail* in MIA PaCa-2 and *Snail, Twist* and, to a degree, *Slug* in PANC-1 is downregulated when cells were treated with either inhibitor (**Supplementary Figure 5B-C**). Consistent with the reduction in cell migration and the expression of EMT markers, actin cytoskeleton staining in compressed cells treated with either inhibitor showed a reduction in cell protrusions as well as stress fibre formation (**Figure 4A-B**). Collectively, these results suggest that mechanical stress can lead to activation of p38 MAPK and JNK signaling pathways to promote cancer cell migration, actin cytoskeleton remodelling and proliferation under compression.

**Figure 3.**
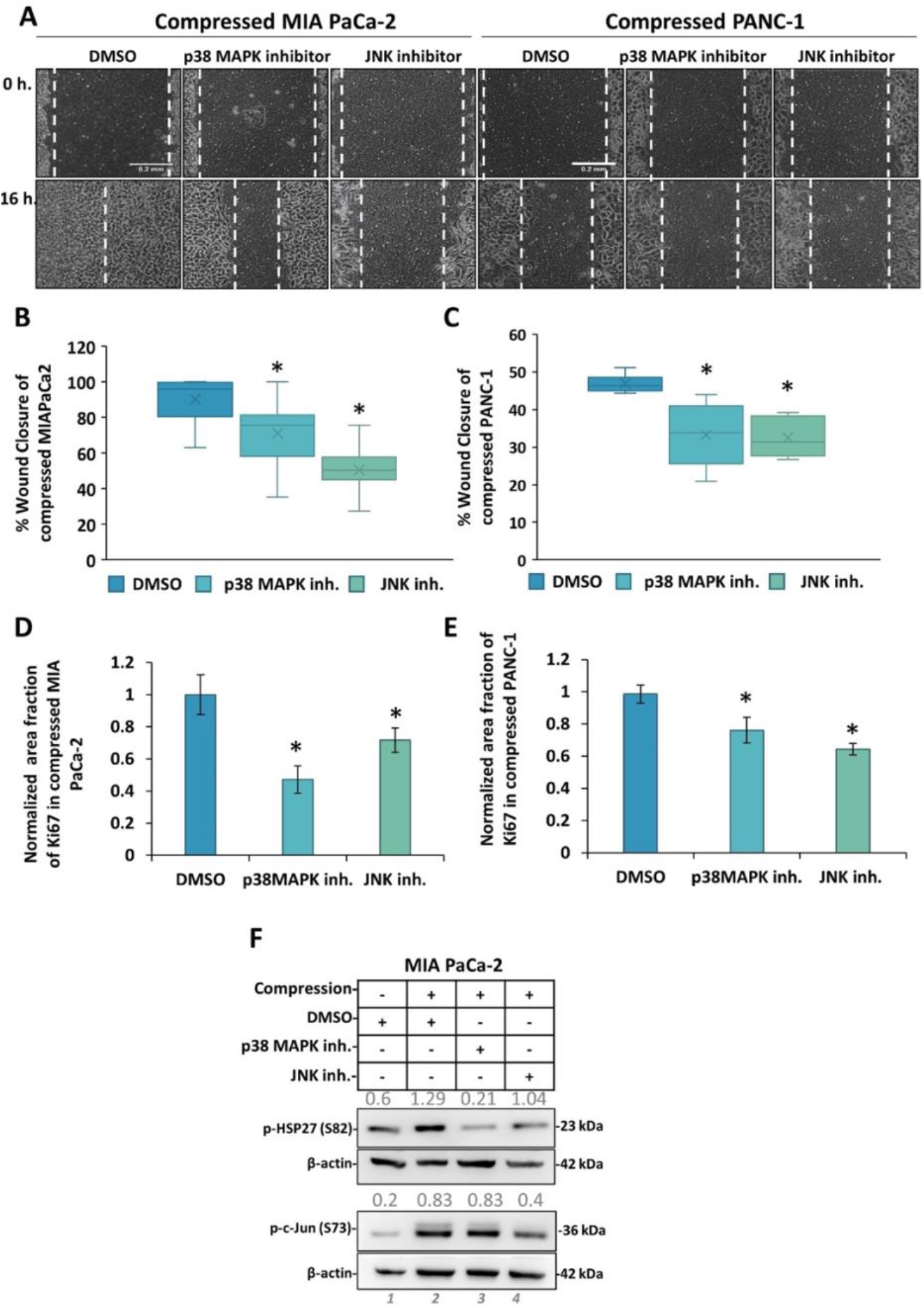
The p38 MAPK/HSP27 and JNK/c-Jun pathways are necessary for both proliferation and migration of pancreatic cancer cells under mechanical stress. **A,** MIA PaCa-2 and PANC-1 cells were pre-treated with 15 μM p38 MAPK inhibitor (SB202190), JNK inhibitor (SP600125) or equal volume of DMSO and subjected to a scratch wound healing assay under 4.0 mmHg of compression. Pictures from at least 4 different fields were taken from three biological replicates. Scale bar: 0.2 mm. White dashed line shows the difference in wound closure between 0 and 16 hours. **B-C,** Graph showing the average ±SE percentage wound closure of compressed MIA PaCa-2 (B) and PANC-1 (C) treated with DMSO or inhibitors from 3 biological replicates (n≥12). Asterisk (*) indicates a statistically significant difference (p<0.05 in student’s *t* test). **D-E**, Graph showing the average Ki67 area fraction in compressed MIA PaCa-2 (D) and PANC-1 (E) cells treated with DMSO, p38 MAPK or JNK inhibitor from at least 10 different fields/ condition from two biological replicates as quantified automatically using an in-house code in MATLAB. Asterisk (*) indicates a statistically significant difference in student’s *t* test (p<0.05). **F**, Representative Western Blotting showing phosphorylated HSP27 (Ser 82) and c-Jun (S73) in control and compressed MIA PaCa-2 cells treated with 15 μM of each inhibitor or equal volume of DMSO. Antibody against β-actin was used as a loading control. Quantification of each antibody compared to loading control was quantified by ImageJ and it is indicated by numbers in grey font.

**Figure 4.**
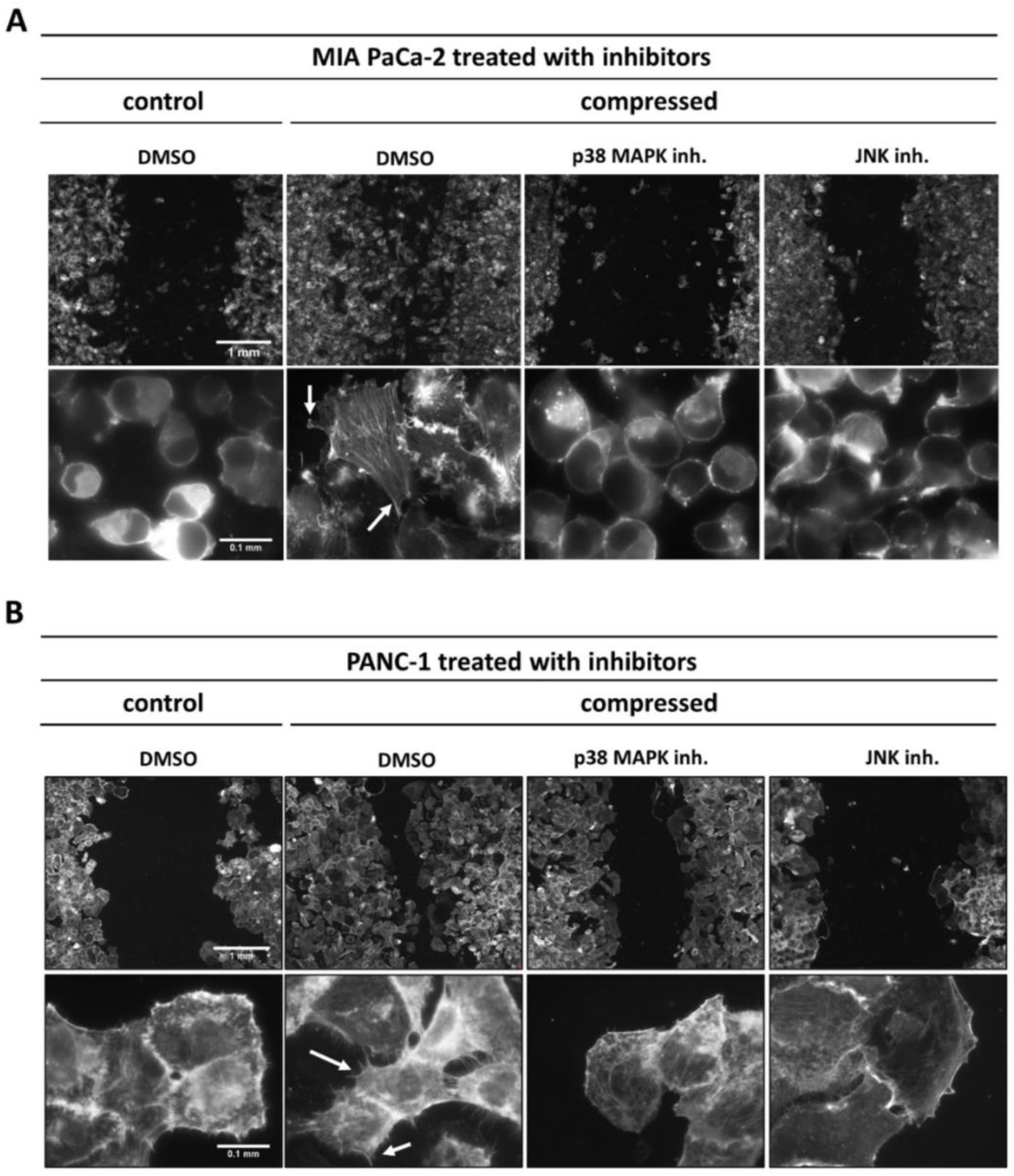
p38 MAPK/ HSP27 and JNK/c-Jun signaling axes promote mechanical stress-induced actin cytoskeleton remodeling. Representative images of phalloidin staining in control and compressed MIA PaCa-2 (A) and PANC-1 (B) cells treated with DMSO, p38 MAPK or JNK inhibitors. In each panel, the first row shows representative images at the end of the wound healing assay, while the second row shows cells at the wound edge in higher magnification (scale bar: 1mm and 0.1mm respectively). White arrows indicate cell protrusions (lamellipodia, filopodia) that are necessary for cell migration.

### The Rac1- and cdc42-GTPases mediate mechanical stress-induced cytoskeletal changes that promote pancreatic cancer cell migration

Since mechanical stress induces the formation of cell protrusions, filopodia and lamellipodia (**Figure 2A-B**), and because these structures are regulated by Rac-1 and cdc42-small GTPases^28–30^, we treated MIA PaCa-2 and PANC-1 cells with siRNAs against *Rac1* and *cdc42* and performed a scratch wound healing assay and phalloidin staining to examine their role in mediating effects of compression. Once we confirmed the successful knockdown of these genes (**Figure 5A-B**), we observed a strong inhibition of wound closure in siRac-1-treated cells, and a weaker effect in the sicdc42-treated cells (**Figure 5C-E**). Consistent with the reduction in migratory potential, knockdown of *Rac-1,* and to a lesser extent of *cdc42,* reduced actin stress fibre, filopodia and lamellipodia formation (**Figure 6A-B, Supplementary Figure 6A**). In line with this, we also found that although Myosin II was activated in either siRNA-treated compressed cells, there was a reduction in actomyosin formation, evidenced by the reduced Myosin II-actin co-localization (**Figure 6**). To determine epistasis between *Rac1* and *cdc42* downregulation and JNK or p38 MAPK activation, we performed Western Blotting. We found that JNK and p38 MAPK activation remained unaffected upon knockdown of either *Rac1* or *cdc42* (**Supplementary Figure 6C**). In addition, treatment with either siRac1 or sicdc42 did not affect the proliferation of either MIA PaCa-2 or PANC-1 cells as indicated by Ki67 staining (**Figure 5F, Supplementary Figure 6B**). Collectively our results suggest that p38 MAPK and JNK activation in response to mechanical stress that sustains cell viability does not depend on the Rac1 and cdc42 GTPases, while these GTPases are necessary for actin cytoskeleton reorganization and cell motility.

**Figure 5.**
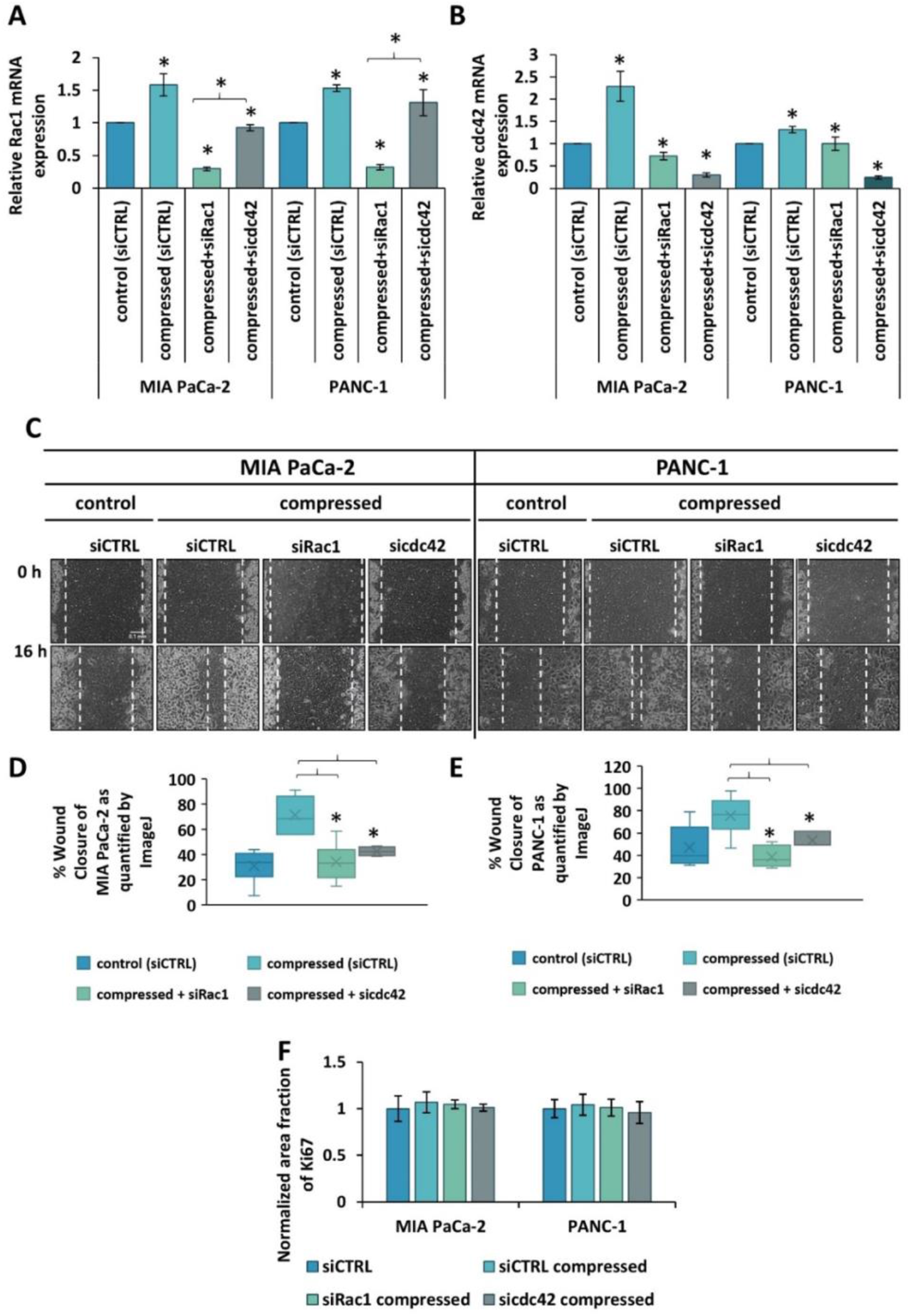
The Rac-1- and cdc42-small GTPases mediate mechanical stress-induced pancreatic cancer cell migration. **A-B**, *Rac1* (A) and *cdc42* (B) mRNA expression was quantified by qPCR in both MIA PaCa-2 and PANC-1. Each bar indicates the mean fold change ±SE of two biological replicates (n=6). Asterisk (*) indicates a statistically significant difference (p<0.05 in student’s *t* test). **C**, MIA PaCa-2 and PANC-1 pancreatic cancer cells were treated with siRNA against *Rac1* (siRac1) *or cdc42* (sicdc42) and then subjected to a scratch wound healing assay for 16 hours under 0.0 or 4.0 mmHg of compression in 2 % FBS containing DMEM. Control cells were treated with stealth siRNA (siCTRL). Scale bar: 0.1 mm. **D-E**, Graphs showing the percentage ±SE wound closure of MIA PaCa-2 (D) and PANC-1 (E) as quantified using ImageJ software. Statistically significant difference in wound closure of compressed siRac1- or sicdc42-treated cells compared to siCTRL-treated cells is indicated with an asterisk (*) (2 biological replicates; n≥6; p<0.05 in student’s *t* test). **F**, Graph showing the average ±SE Ki67 area fraction in MIA PaCa-2 and PANC-1 control and compressed cells treated with siCTRL, siRac1 or sicdc42 from at least 5 different fields/ condition from two biological replicates as quantified automatically using an in-house code in MATLAB. No statistically significant changes were observed.

**Figure 6.**
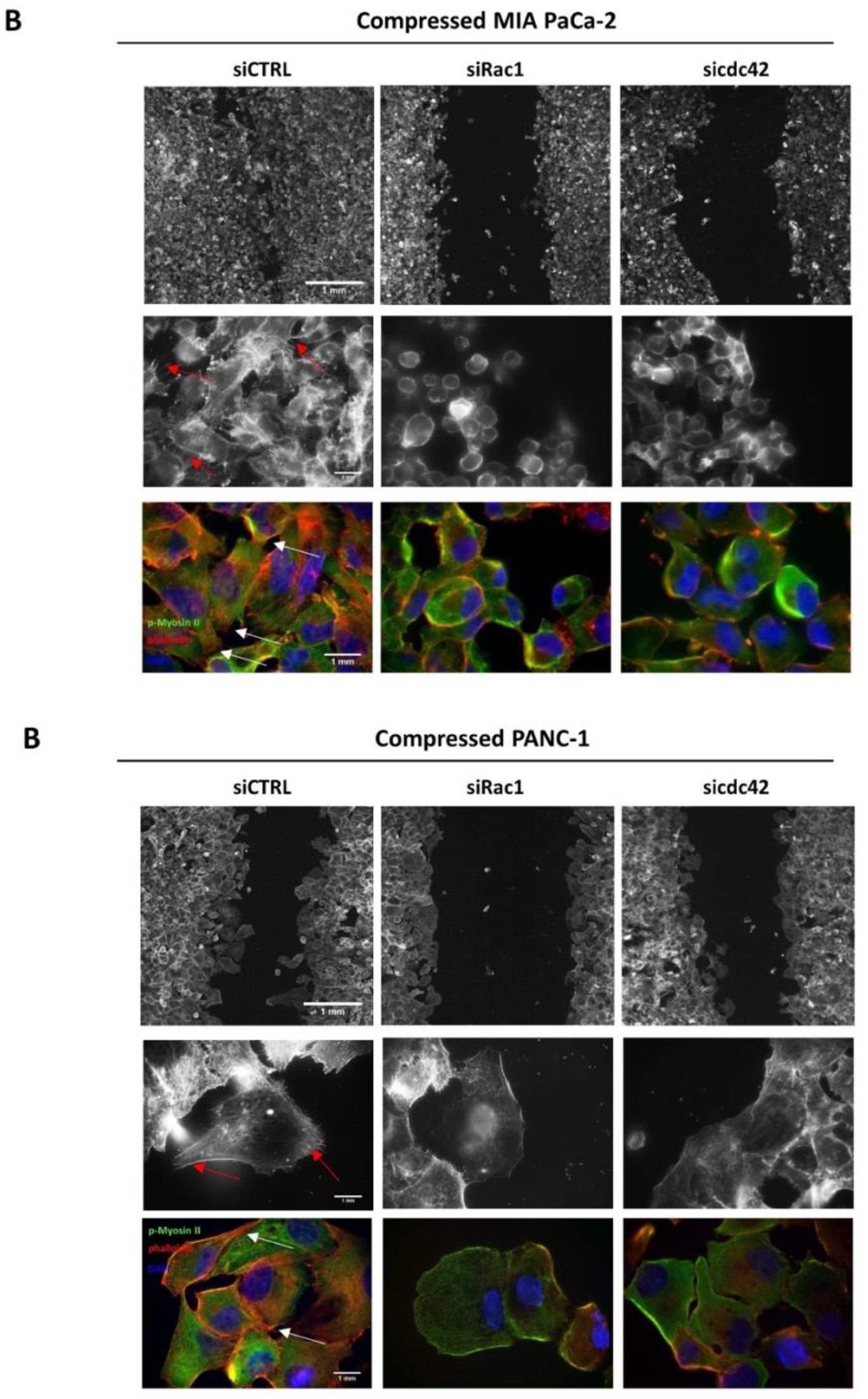
Rac-1 and cdc42 small GTPases mediate actin cytoskeleton remodelling and contractility to induce cell migration under stress. Representative images of phalloidin and phospho-Myosin II (p-Myosin II) staining in compressed MIA PaCa-2 (A) and PANC-1 (B) cells treated with siCTRL, siRac-1 or sicdc42. In each panel, the first row shows representative images at the end of the wound healing assay, while the second row shows cells at the wound edge in higher magnification (scale bar: 1mm). Red arrows indicate cell protrusions that are necessary for cell migration. Third row shows p-Myosin II and phalloidin staining in high magnification (scale bar: 1mm).

## Discussion

Although mechanical stress has already been associated with pancreatic tumour progression and treatment resistance, little is known about the molecular mechanisms involved. To this end, we used RPPA-based proteomic profiling, in order to examine signaling pathway adaptation in compressed pancreatic cancer cells; we evaluated multiple signaling pathways, such as actin cytoskeleton remodelling, Ras/MAPK, PI3K/ Akt, TSC/mTOR, cell cycle progression and apoptosis. Based on RPPA, we observed an upregulation of the autophagy marker LC3A-B as well as the ER stress sensor, BiP-GRP78. This could be linked with the observation that mechanical compression has recently been suggested to cause mitochondrial dysfunction, leading to autophagy and ER stress response activation through alterations in actin cytoskeleton organization and dynamics^31,32^. Moreover, it has been suggested that both the autophagy and ER stress response could be mediated through p38 MAPK and JNK signaling pathway activation, which are indeed activated in compressed cells^31,32^. Thus, it is possible that mechanical stress can induce cytoskeletal changes, and signaling pathway adaptation that leads to autophagy and ER stress response, which, in turn, promote cell growth and survival under environmental stress. This is also supported by data showing that autophagy could mediate the Ras-driven pancreatic tumour progression and could serve as a marker of poor prognosis for pancreatic tumour patients^33^. Notably, the mechanical stress, induced activation of autophagy and ER stress response could also explain that while targets of the mTOR and cell cycle progression were downregulated or inactivated in compressed cells, the proliferation of compressed cells did not exhibit significant changes compared to uncompressed cells at least 16 hours post-compression (**Supplementary Figure 4**).

The most pronounced changes found in RPPA analysis of compressed cells were related to stress response, cell migration and actin cytoskeleton organization (**Figure 1**). To this end, and in order to identify the mechanical stress-induced mechanism that caused those alterations, we firstly focused on p38 MAPK and its downstream target HSP27, as well as the JNK and its downstream target c-Jun exhibited activation in compressed cells. Although it has already been shown that increased levels of p38 MAPK and JNK activation enhance the efficacy of chemotherapeutic agents, increasing patient survival^19,34^, it is found that both kinases could also lead to increased cancer cell growth, migration and invasion^19, 35, 36^. On a molecular basis, this could depend on the different levels of kinase activity, the duration of its activation, the interaction of the kinase with other signaling pathways as well as cell-type differences. The role of p38 MAPK in cell migration, could be mediated by its downstream target, HSP27, which has been previously shown to promote growth, migration, actin cytoskeleton remodelling and resistance to chemotherapy in pancreatic tumours and is associated with poor survival^37,38^ for patients with pancreatic cancer^37,39,19,27^. Similar to p38 MAPK, JNK has also been implicated in cancer cell migration as it has been shown to phosphorylate paxillin to induce the migration and invasion of pancreatic cancer cells^40, 41^, while its main downstream target, c-Jun, is expressed in 87% of pancreatic tumour lesions and has been implicated in pancreatic tumour progression^42^. To this end, and to further explore whether these molecules are responsible for the increased migratory ability of compressed cells, we first treated MIA PaCa-2 and PANC1 cells with inhibitors for the p38 MAPK and JNK signaling cascades and found that both are necessary for cell migration and growth under compression (**Figure 3–4**). Their role in cell growth could partially be supported by their contribution in mediating autophagy and ER stress response, events that are necessary for cell adaptation under compression. Regarding their role in cell migration, we further showed that both p38 MAPK and JNK can regulate expression of EMT genes and are necessary for actin cytoskeleton remodelling (**Figures 3–4**). Interestingly, while HSP27 is a downstream target of p38 MAPK that regulates the organization of actin filaments, the disruption of actin cytoskeleton in compressed cells was observed in cells treated with either JNK or p38 MAPK inhibitor. This could be explained by impaired HSP27 activation following JNK inhibition (**Figure 3, Supplementary Figure 5**). Although these kinases have been previously reported to have antagonistic effects in cell proliferation and survival ^19^, it is important to consider the tumor type, as well as the biomechanical cues present in the tumor microenvironment, prior to their targeting for cancer therapy.

Next, based on the increased formation of cell protrusions and Myosin II activation observed in compressed cells (**Figure 1–2**), and because Rac-1 and cdc42 small GTPases showed an activation in G-LISA and are known upstream activators of Myosin II to facilitate cell motility^43–45^, we downregulated *RAC1* and *cdc42* using siRNAs to examine their role in the mechanical stress-induced migration. Phalloidin and phospho-Myosin II staining in compressed siRAC1- and sicdc42-treated cells confirmed the role of RAC1 and cdc42 in the formation of cell protrusions and actomyosin contractility, while a scratch wound healing assay revealed that they are all necessary for cell migration under mechanical stress conditions. This effect was less pronounced in sicdc42-treated cells, which exhibited more cell protrusions compared to siRac1-treated cells (**Supplementary Figure 6A**). This could be explained by the observation that in sicdc42-treated cells, *Rac1* mRNA expression was higher compared to siRac1-treated cells, suggesting that Rac1 could have a dominant role in the formation of cell protrusions.

Collectively, our results, establish a novel mechanism by which mechanical stress enhances tumor progression through the activation of p38 MAPK/HSP27 and JNK/c-Jun signaling axes, and actin cytoskeleton remodelers, Rac1, cdc42, and Myosin II (conceptual model shown in **Figure 7**). Moreover, our proteomic profiling data suggest that cancer cells could survive under mechanical stress conditions through activation of autophagy and ER stress response, indicated by the upregulation of LC3A-B and GRP78. The high expression of these molecules in pancreatic tumors could be eventually used as a novel marker for the presence of mechanical forces in the tumor interior, and further enhance the importance of targeting mechanically-triggered signaling, in combination with conventional treatments, for the cure of pancreatic tumor patients. Therapeutic strategies to alleviate mechanical stresses in tumors have already been developed, with the aim to overcome barriers of effective drug delivery into the highly dense microenvironment of pancreatic and other desmoplastic tumors^46^. Our results highlight that targeting aberrant signaling in tumor cells adapting to the mechanical tumor microenvironment presents a novel approach to block tumor migration.

**Figure 7.**
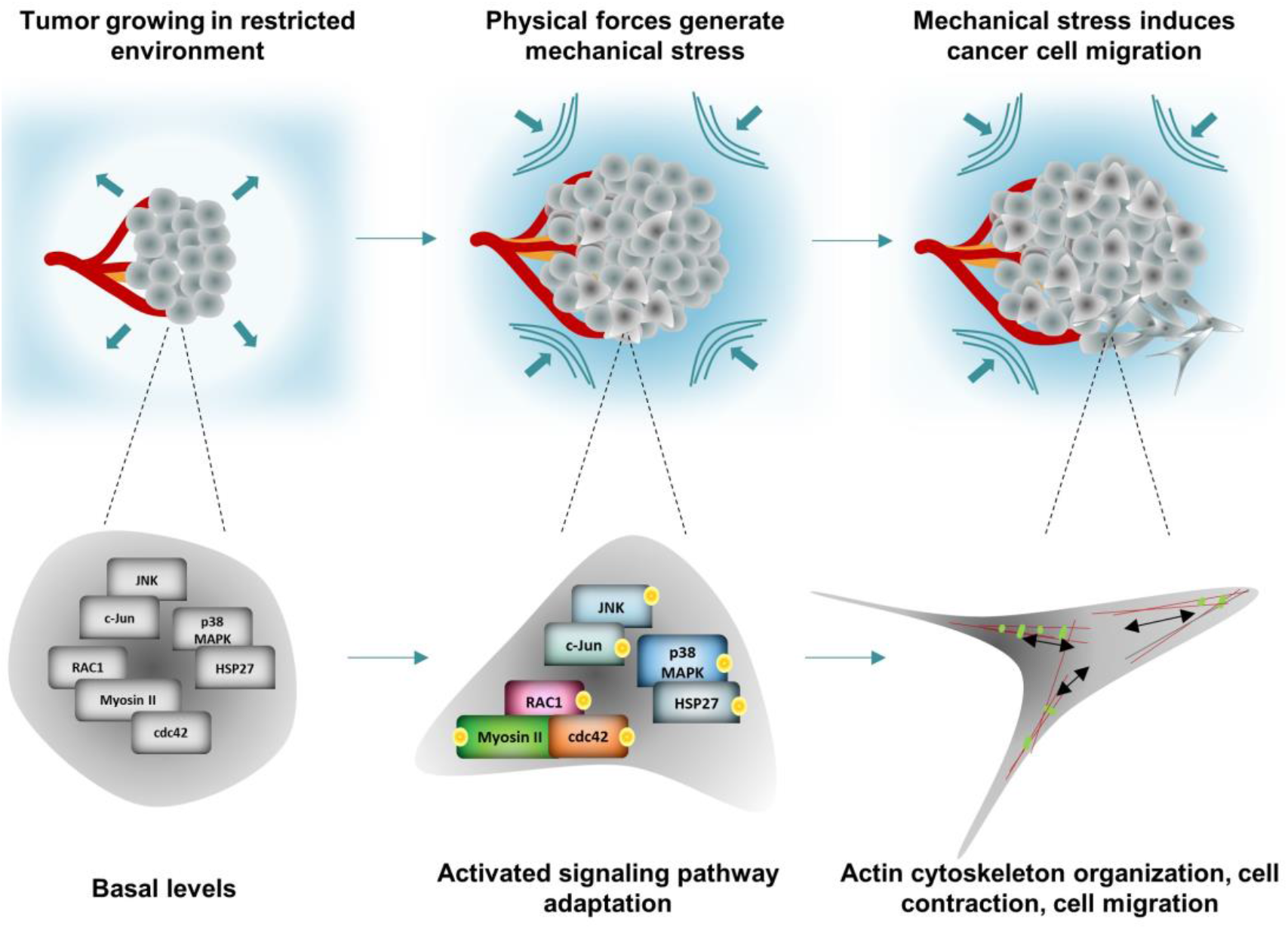
Mechanical stress-induced signaling pathway adaptation that drives pancreatic cancer cell migration. Mechanical compressive forces are generated during pancreatic tumor growth in the restricted environment of a host tissue. These forces transmit the respective solid stress intracellularly, by activation of JNK/c-Jun, p38 MAPK/HSP27, Rac1 and cdc42. The activation of HSP27 along with activated Rac1 and cdc42 in turn regulate actin cytoskeleton remodeling for the formation of cell protrusions changing progressively the cell shape. Rac1 and cdc42 mediate actomyosin contractility, to eventually promote pancreatic cancer cell migration under compression.

## Materials and Methods

### Antibodies

Antibodies against phospho-Heat Shock Protein 27 (S82) and phospho-c-Jun (S73) were obtained from Abcam (Cambridge, United Kingdom), while phospho-JNK (T183/Y185) and phospho-p38 MAPK (T180/Y182), were obtained from Cell Signaling Technology (Danvers, MA, USA). Antibody against β-actin was obtained from Sigma-Aldrich (St. Louis, MO, USA) and anti-β-tubulin from Developmental Studies Hybridoma Bank (DSHB). For the immunofluorescence staining, phalloidin was obtained from Biotium (California, United States), phosphor-Myosin II (Ser1943) from Cell Signaling Technology (Danvers, MA, USA) and Ki67 from Abcam (Cambridge, United Kingdom).

### Cell Lines

MIA PaCa-2 and PANC-1 were purchased from the American Type Culture Collection (ATCC, Manassas, VA, USA). Cells were grown in Dulbecco’s Modified Eagle Medium supplemented with 10% Fetal Bovine Serum (FBS), 1% Antibiotic/Antimycotic, and incubated in a CO_2_-incubator at 37°C.

### Application of mechanical stress

For the application of a defined and controlled compression (4.0 mmHg) on cancer cells, a previously described transmembrane pressure device was employed ^20, 47^. 24 hours prior to compression, the culture medium switched to 2% FBS-containing medium. A piston with adjustable weight was added on the top of a cell monolayer covered with an agarose cushion and placed in a transwell insert, in order to apply 4.0 mmHg of compressive stress for 16 hours. Control cells were covered with an agarose cushion only.

### Protein Extraction for Reverse Phase Protein Array (RPPA)

MIA PaCa-2 pancreatic cancer cells were compressed by 0.0 or 4.0 mmHg in 2% FBS-containing medium for 16 hours and washed twice with cold PBS. Cells were then lysed using 150μl of cold lysis buffer containing 50mM HEPES, 150mM NaCl, 1.5mM MgCl2, 1mM EGTA, 10% Glycerol and freshly added 1X Halt Protease Inhibitor Cocktail (Thermo Fischer Scientific), 1mM Na3VO4, 100mM NaF, 10mM Sodium Pyrophosphate and 1% Triton-X. Cell lysates were then centrifuged at 14000 rpm for 15 minutes at 4°C, and supernatants were collected and mixed with 4X sample buffer containing 40% Glycerol, 8% SDS, 0.25M Tris-HCl pH 6, 10% beta-mercaptoethanol (3:1). Finally, lysates were boiled at 95°C for 5 minutes and kept at −80°C. Proteomic analysis using RPPA was performed at the RPPA core facility (MD Anderson Cancer Center, Houston, TX).

### RPPA data analysis

For the RPPA analysis, pairwise t-test was performed to identify proteins that are differentially expressed between compressed and uncompressed cells. Differences are considered significant when p < 0.05. To visualize the protein expression patterns of compressed and uncompressed MIA PaCa-2 cells, heatmaps were generated with the *gplots* R-package. Proteins that showed significance in the paired t-test were included in the heatmap. Heatmaps included proteins sorted based on their fold-change (compressed/control). For *pathway analysis,* scores were used to evaluate the activity of the pathways after compression. Pathway scores were calculated as the average weighted (positive or negative) sum of expression level of all members in each pathway^27, 48^ and normalized to the uncompressed expression level. Members of 9 pathways are showed in **Supplementary Figure 2**. Heatmaps for pathways were generated to show the compression-induced log2-ratio of each protein member.

### Cell treatments

In order to examine the role of p38 MAPK and JNK in the migratory ability of pancreatic cancer cells under 4.0 mmHg stress, MIA PaCa-2 and PANC-1 cells were grown in 2% FBS-containing DMEM for 24 hours and were then pre-treated with 15 μM of p38 MAPK inhibitor (SB202190, MedChemExpress) or JNK inhibitor (SP600125, MedChemExpress) for 1 hour. Control cells were pre-treated with equal volume of solvent (DMSO). The concentration of each inhibitor was selected based on previously published studies employing pancreatic cancer cells^49–51^. Mechanical stress (4.0 mmHg) was then applied on cells growing in 2% FBS-containing medium in the presence of the inhibitors for 16 hours.

### Transient transfection of pancreatic cancer cells with siRNAs against Rac1 and cdc42

To determine whether the Rac1 or cdc42 small GTPases are implicated in the mechanical migration of pancreatic cancer cells, cells were treated with stealth siRNA (negative control, Invitrogen), siRac1 or sicdc42 (Invitrogen) using Lipofectamine 2000 transfection reagent (Invitrogen) for 24 hours according to the manufacturer’s guidelines. Cells were then washed twice with PBS and medium was switched to 2% FBS-containing medium for another 24 hours. Finally, mechanical stress was applied as described above for 16 hours.

### RNA Isolation and Real-Time PCR

Total RNA was extracted from cancer cells as previously described^47^. Briefly, RNA extraction was accomplished using Trizol (Invitrogen). RNA was then transcribed to cDNA using Superscript III Reverse Transcriptase (Invitrogen) and cDNA (1:10) was employed for quantification of gene expression by real-time PCR using β-actin as a reference gene. All primers used are shown in **Supplementary Table 1**. Quantification of relative gene expression was performed using the ΔΔCt method, where data were log transformed and analyzed using standard methodology.

### Protein Extraction and Western Blot Analysis and Quantification

For protein expression analysis, a standard immunoblotting protocol was followed as described previously^20, 21^. Western Blotting images were analyzed in Adobe Photoshop. Images were converted to grayscale. In some cases, contrast was used in the entire blot to enhance image clarity. No other image manipulation was performed. Uncropped images of Western Blotting used in this article can be found in **Supplementary Material**.

### Wound Healing

A wound healing assay was performed on cancer cells according to previous studies^20^. Images from at least 4 different fields per condition were taken at 0 hours and 16 hours. The cell-free area was quantified using the ImageJ software. Quantification was performed for each condition using the following formula:

(Width of the wound at 0 hours – Width of the wound at 24 hours) / (Width of the wound at 0 hours).

### Immunofluorescence Staining

To determine the effect of mechanical stress on cell shape and proliferation, cells were fixed in 4% paraformaldehyde, permeabilized in PBS containing 0.25 % Triton X-100 and blocked with PBS containing 0.1 % Tween-20 and 1 % BSA. Cells were then stained with phalloidin or anti-Ki67 antibody diluted in blocking buffer for 30 minutes or 1 hour respectively, at room temperature. Alexa 488-goat anti-rabbit antibody was used as a secondary antibody for anti-Ki67. Images were obtained using a fluorescent microscope (Olympus BX53). For image analysis, the calculation of Ki67 area fraction was performed automatically using a previously developed inhouse code in MATLAB (MathWorks, Inc., Natick,MA) ^13, 14, 52^.

### G-LISA assays

To examine whether Rac1 and cdc42 small GTPases are activated in response to mechanical stress, the Rac1- and cdc42 Small GTPase Activation Assay kits (G-LISA) were used (027BK127-S, 027BK128-S, Cytoskeleton, Inc) following the company’s guidelines.

### Statistical Analysis

Results are represented as mean ± standard error (SE). Significant changes were determined by Student’s *t*-test using two-tail distribution. Differences with *p*-values <0.05 were considered as significant (indicated by an asterisk *).

### Data Availability

RPPA data are available via this link: http://dx.doi.org/10.17632/b3mydj4grf.1

## Author Contributions

MK was responsible for designing the study, performing the *in vitro* experiments, analyzing data, interpreting results, and writing the paper; RL was responsible for analyzing the data of RPPA and writing the paper; GBM contributed to the RPPA assay, interpreting results and writing the paper; TS was responsible for designing the study, supervising experiments, interpreting results and editing the manuscript; IZ was responsible for supervising the study, supervising RPPA data analysis, interpreting results and editing the manuscript. All authors have read and approved the final manuscript.

## Appendix I

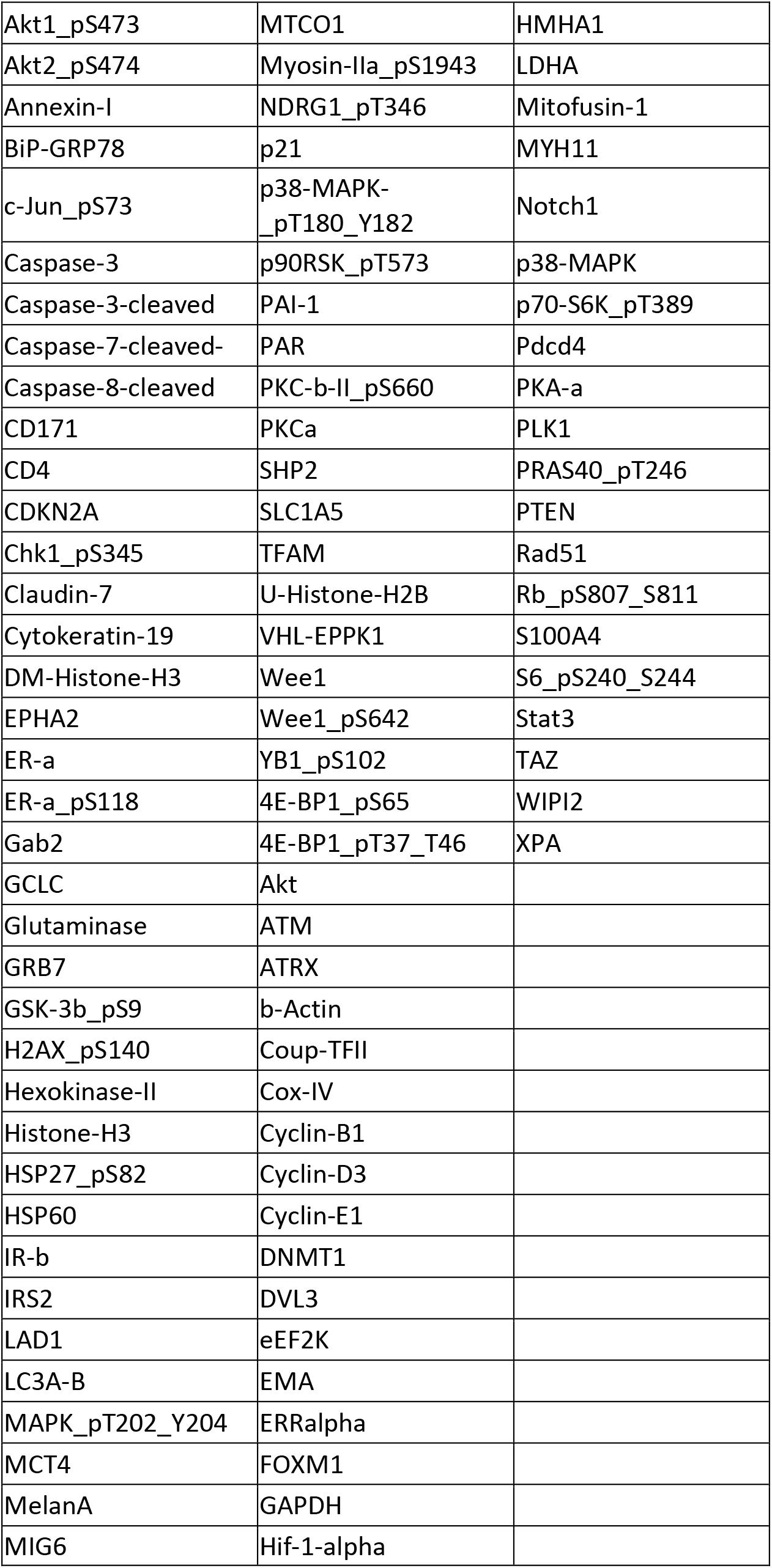
Proteins with >20% foldchange for Figure 1 A

