## Supplementary data for "Mechanical stress in pancreatic cancer: Signaling pathway adaptation activates cytoskeletal remodeling and enhances cell migration"

### SUPPLEMENTARY FIGURES AND TABLES

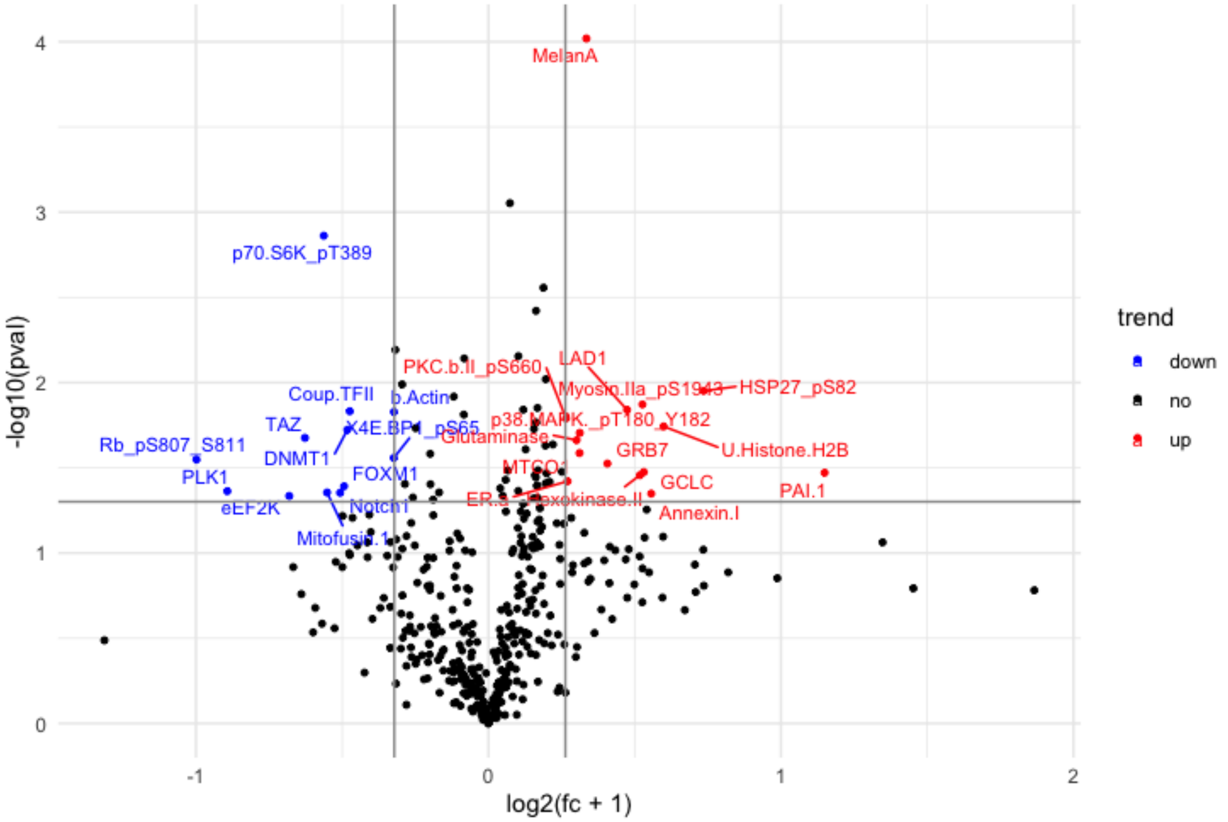


**Supplementary Figure 1. Volcano plot of proteins altered by mechanical stress.** Values represent the log2-ratio (compressed/uncompressed) of expression level for each protein in each condition (n=3; 3 biological replicates). All proteins are depicted in this plot, with black color for no significant change, red for increase (>20%) and blue for decrease (<20%).

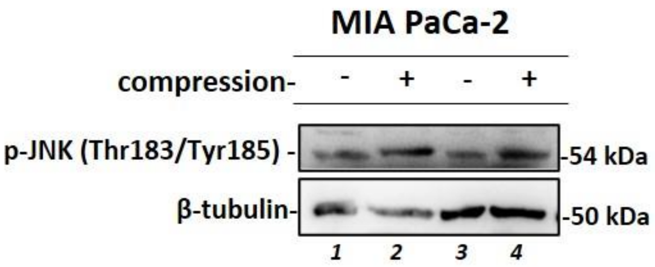


**Supplement Figure 2. Western Blotting showing activated JNK (Thr183/Tyr185) in control and compressed MIA PaCa-2 cells.**

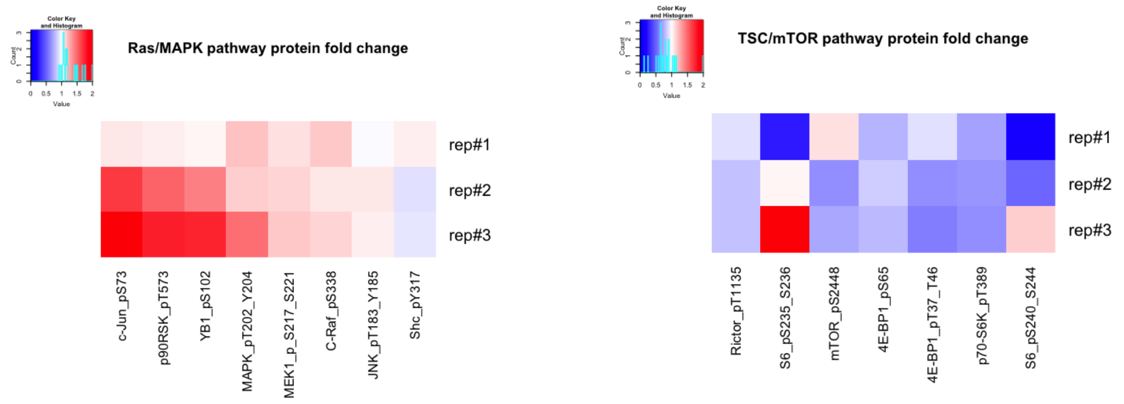


**Supplement Figure 3. Mechanical stress upregulates the RAS/MAPK and downregulates the TSC/mTOR pathway.** Heatmaps showing the expression level of each protein member in the selected pathways. Values in the heatmap represent the log2-ratio (compressed/uncompressed) of expression level for each protein in each condition (n=3; 3 biological replicates). Red: increase vs. blue: decrease

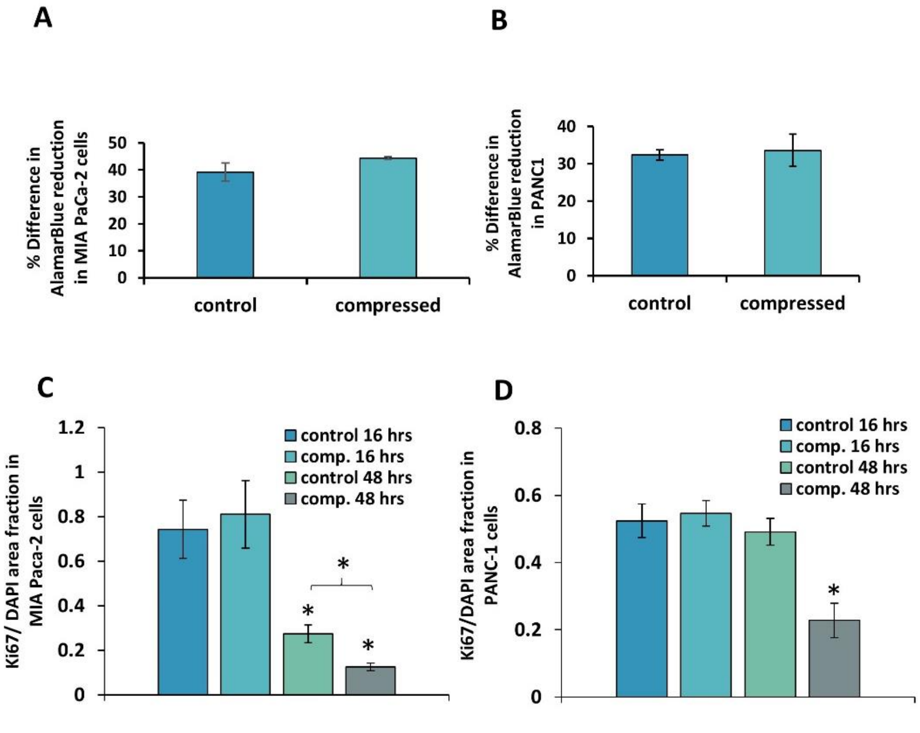


**Supplementary Figure 4. The effect of mechanical stress on the proliferation of pancreatic cancer cells. A-B,** MIA PaCa-2 and PANC-1 cells were subjected to 0.0 (agarose cushion only) or 4.0 mmHg of compressive stress for 16 hours, and AlamarBlue assay was performed prior- and post-compression. The absorbance measured by AlamarBlue reagent represents the metabolic activity of the cells, and thus it was used as an indicator for cell viability. Graphs represent the average % difference ±SE in AlamarBlue reduction prior-and post-compression in control and compressed cells. No statistically significant differences were observed in compressed cells as compared to the control (2 biological replicates; n=6, *p<0.05 in student's t test). **C-D**, MIA PaCa-2 and PANC-

1 cells were compressed for 16 and 48 hours and then were stained with Ki67, a marker for proliferation. The average Ki67 area fraction from at least 8 different fields/condition from two biological replicates was quantified automatically using an in-house code in MATLAB. Asterisk (*) indicates a statistically significant difference ($p<0.05$ in student's t test).

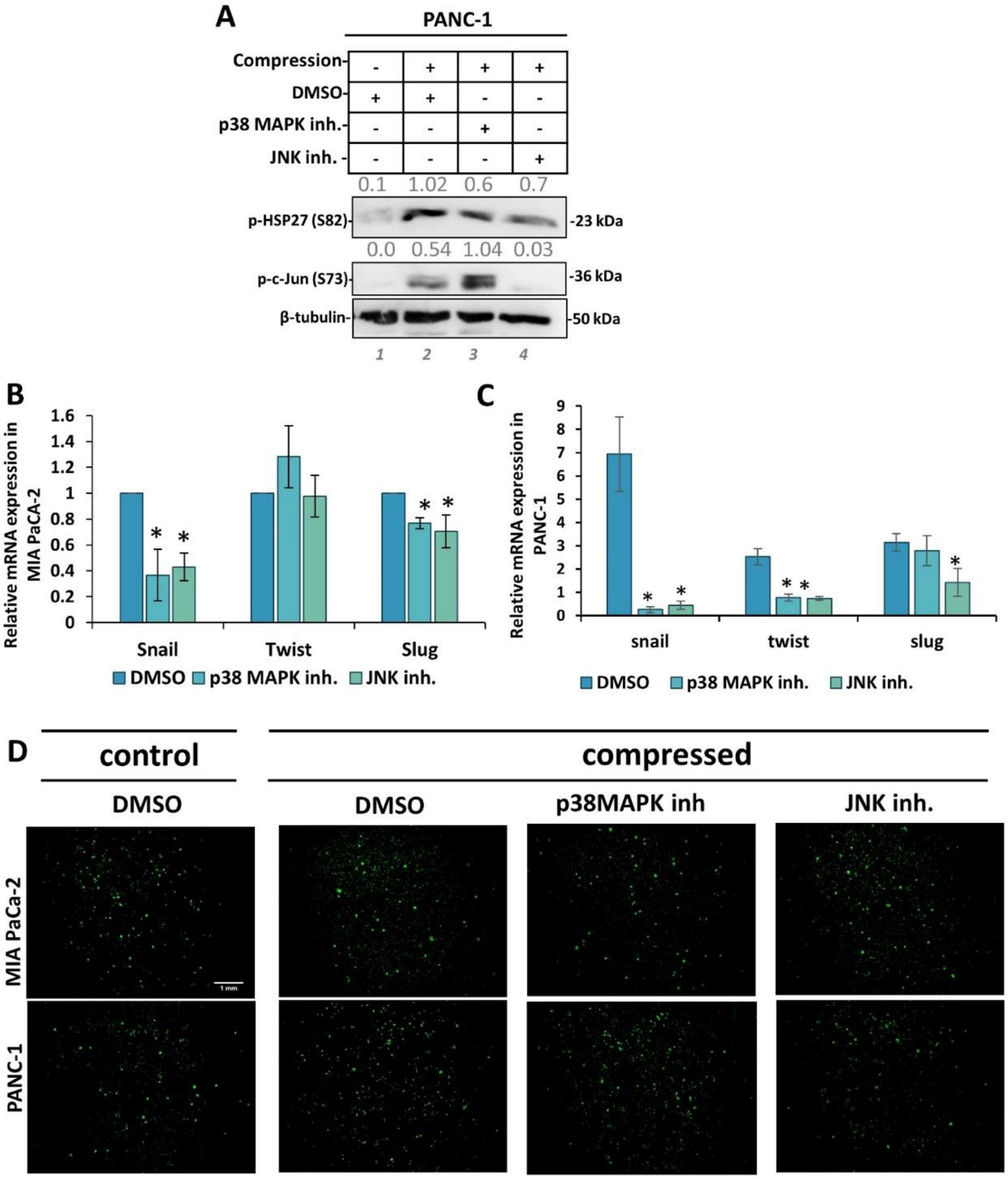


**Supplementary Figure 5 A**, Representative Western Blotting showing phosphorylated HSP27 (Ser 82) and c-Jun (S73) in control and compressed PANC-1 cells treated with 15 μM of each inhibitor or equal volume of DMSO. Antibody against β-actin was used as a loading control. Quantification of each antibody compared to loading control was quantified by ImageJ and it is indicated by numbers in grey font. **B-C**, qPCR was used to quantify the mRNA levels of EMT markers in compressed MIA PaCa-2 (B) and PANC-1 (C) cells treated with 15 μM of each inhibitor compared to compressed cells treated with DMSO. ΔΔCt method was used to quantify the gene expression in each sample using as a reference the expression in compressed and treated with DMSO cells. Bar graphs represent the mean fold change ±SE of two biological replicates experiments (n=6) and statistical changes are indicated with an asterisk (*) ($p<0.05$ in student *t* test). **D**, Representative images of Ki67 staining in MIA PaCa-2 and PANC-1 uncompressed (DMSO) and compressed cells (DMSO+4 mmHg), and compressed cells treated with p38 MAPK or JNK inhibitors. Scale bar: 1mm.

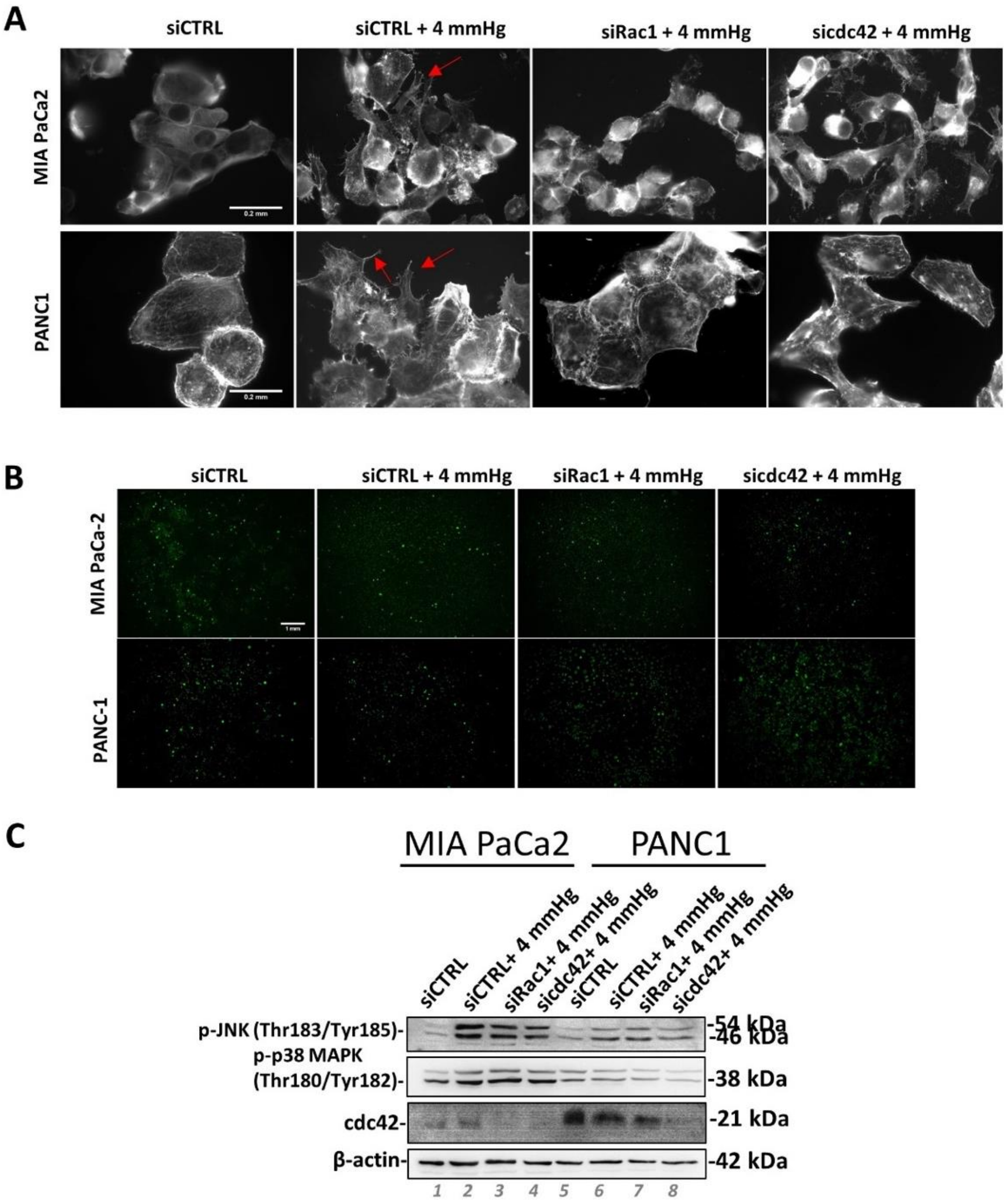


**Supplementary Figure 6. A,** Representative images of phalloidin staining in control and compressed MIA PaCa-2 and PANC-1 cells treated with siCTRL, siRac-1 or sicdc42. Red arrows indicate cell protrusions that facilitate cell migration. Scale bar 0.2 mm. **B,** Representative images of Ki67 staining in uncompressed MIA PaCa-2 and PANC-1 treated with siCTRL and compressed treated with siCTRL, si*Rac* or sicdc42. Scale bar: 1mm. **C**, Representative Western Blotting in control and compressed MIA PaCa-2 and PANC-1 treated with siCTRL, si*Rac1* or sicdc42 for the detection of the phosphorylated levels of JNK (Thr183/Tyr185), p38 MAPK (Thr180/Tyr182) and total levels of cdc42. Antibody against β-actin was used as a loading control.

**Full-length Western Blotting Images.**

**Figure 2A**

**-p-HSP27**

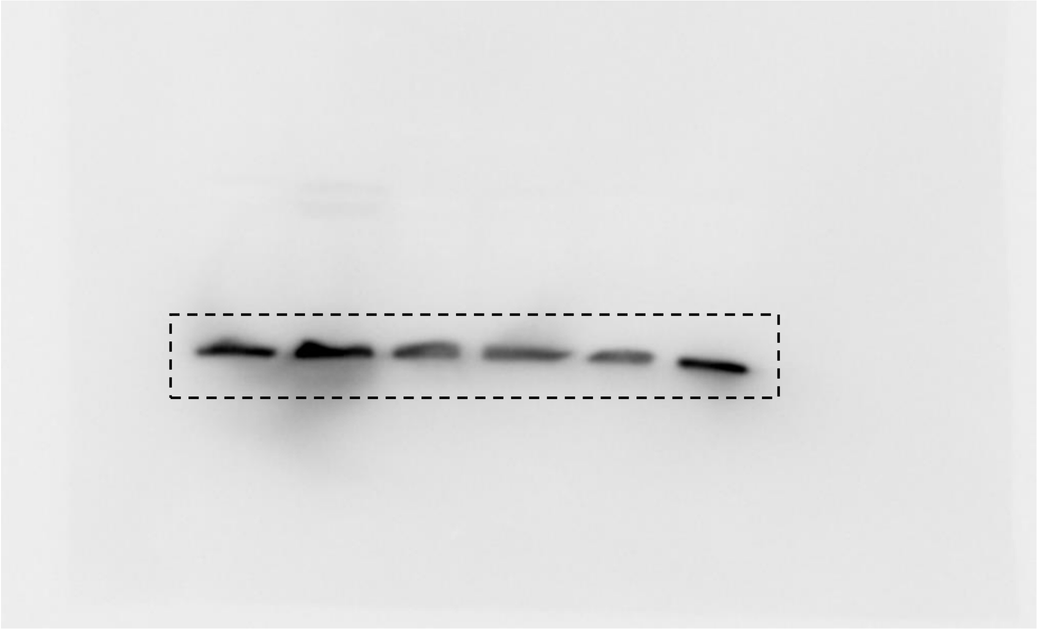

**b-tubulin**

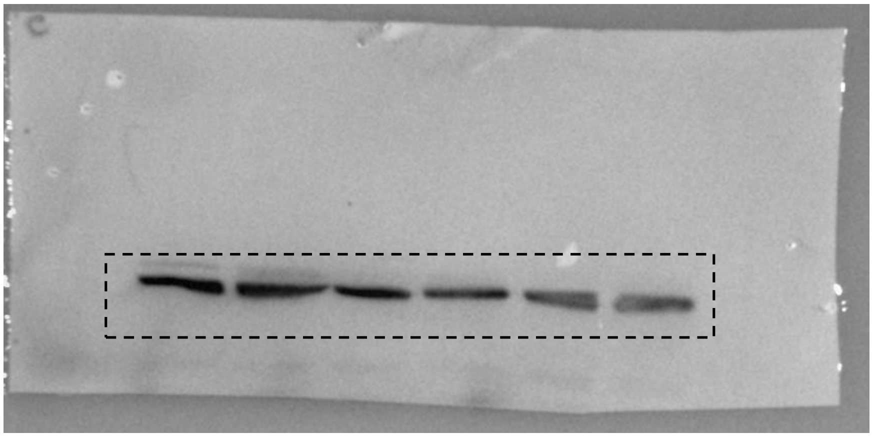

Figure 2C

**p-p38 MAPK**

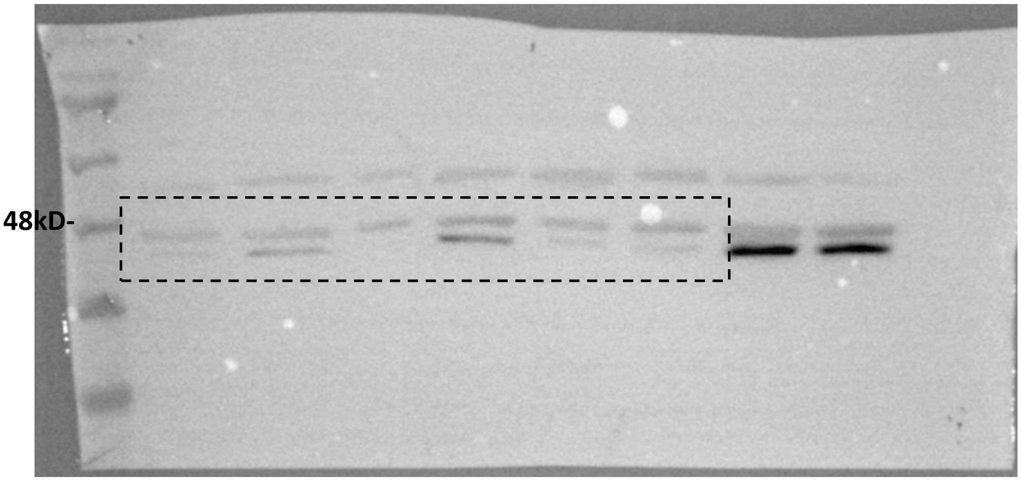


**-b-tubulin**

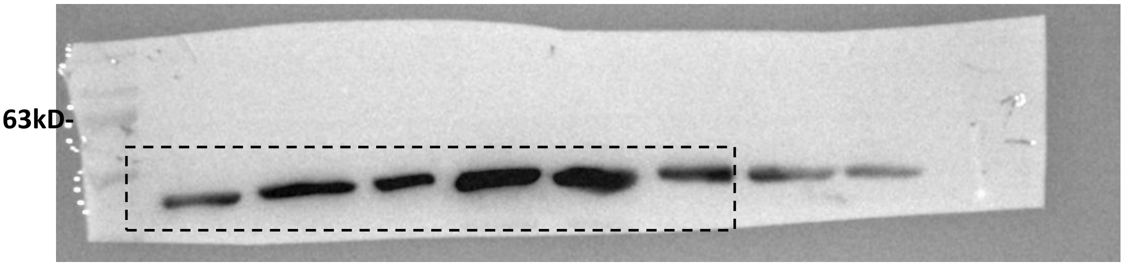


Figure 2E

**-c-Jun**

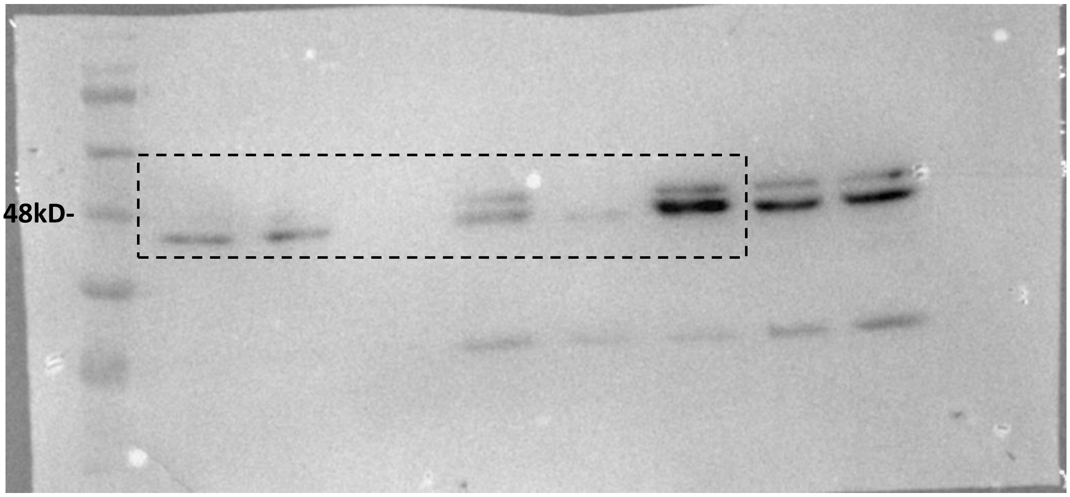


**-b-tubulin**

63kD-

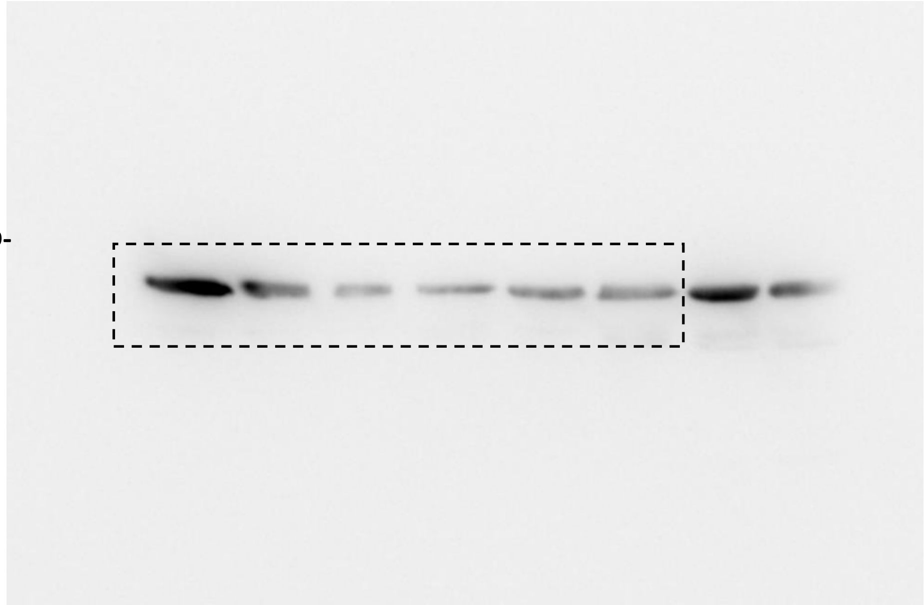

**<u>Figure 2G</u>**

**-pHSP27**

35kD-

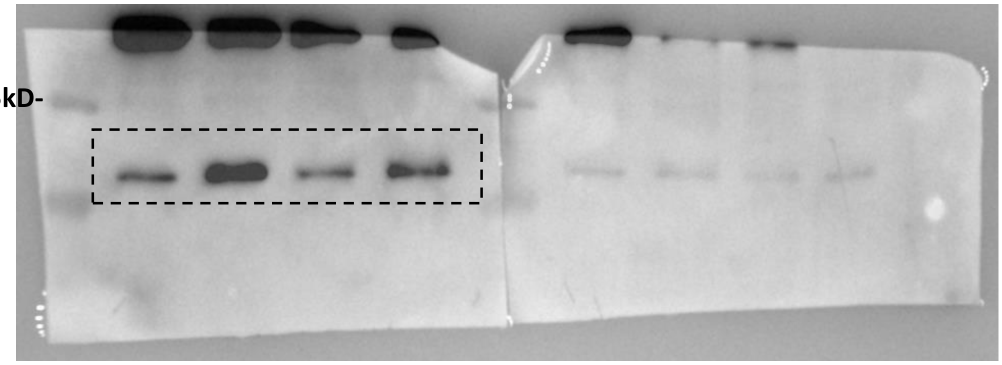

**-pJNK**

48kD-

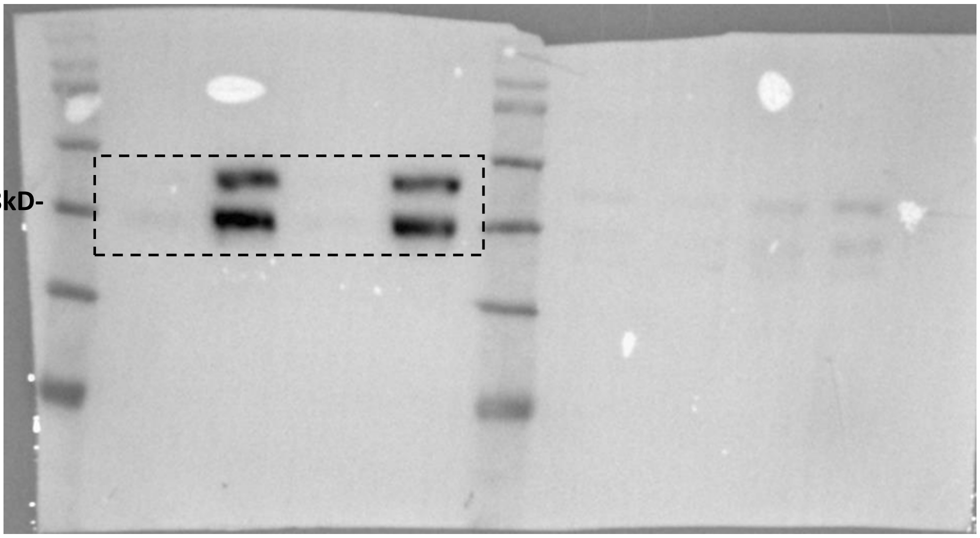

**-p-p38 MAPK**

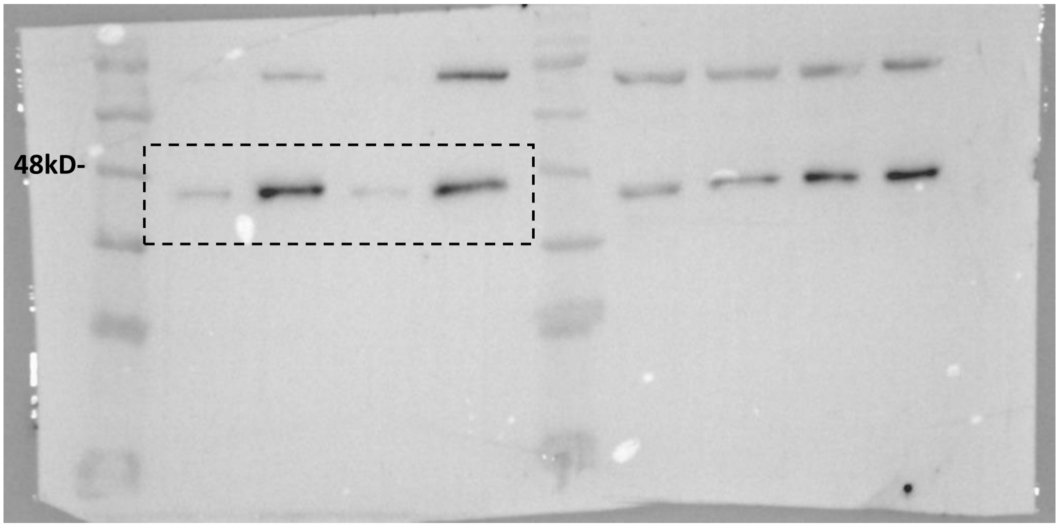


**-b-tubulin**

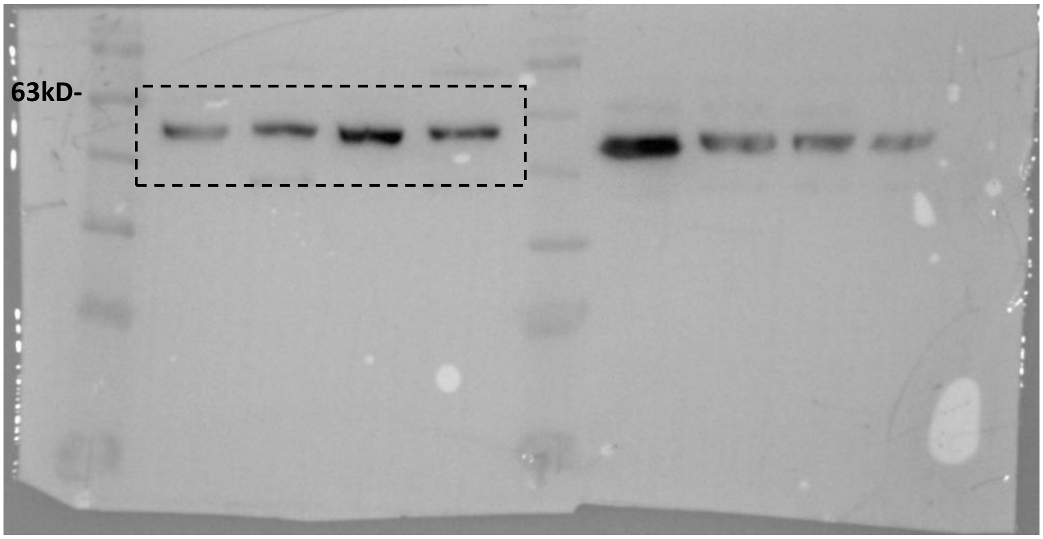

### Supplementary Figure 3

**-pJNK**

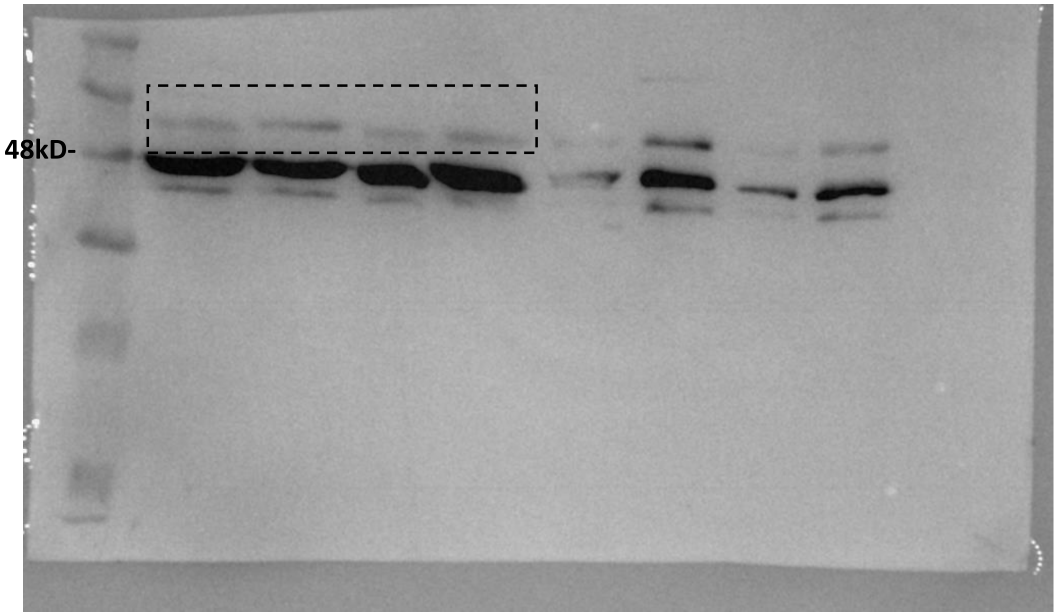


**-B-TUBULIN**

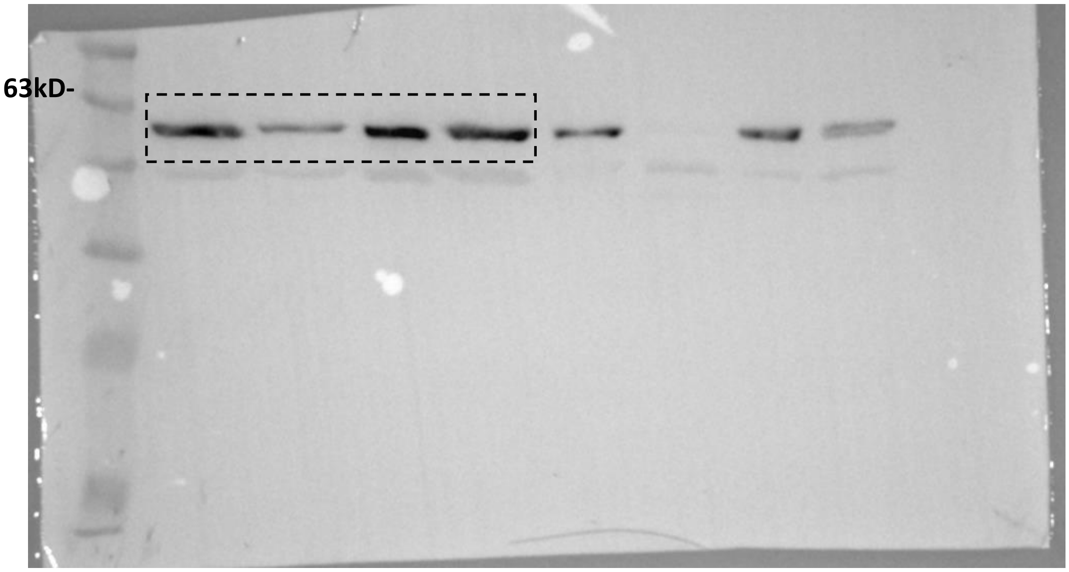

Figure 4F

-pHSP27 -actin on HSP27

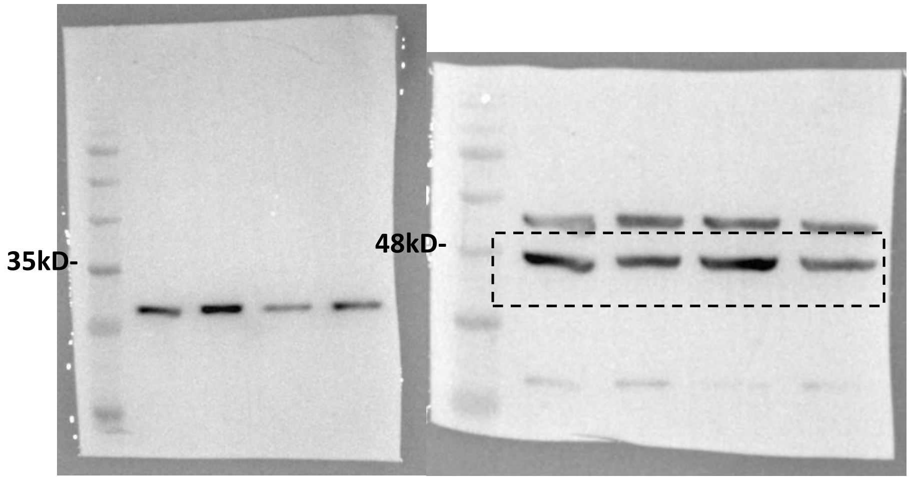


-p-c-Jun Actin on c-Jun

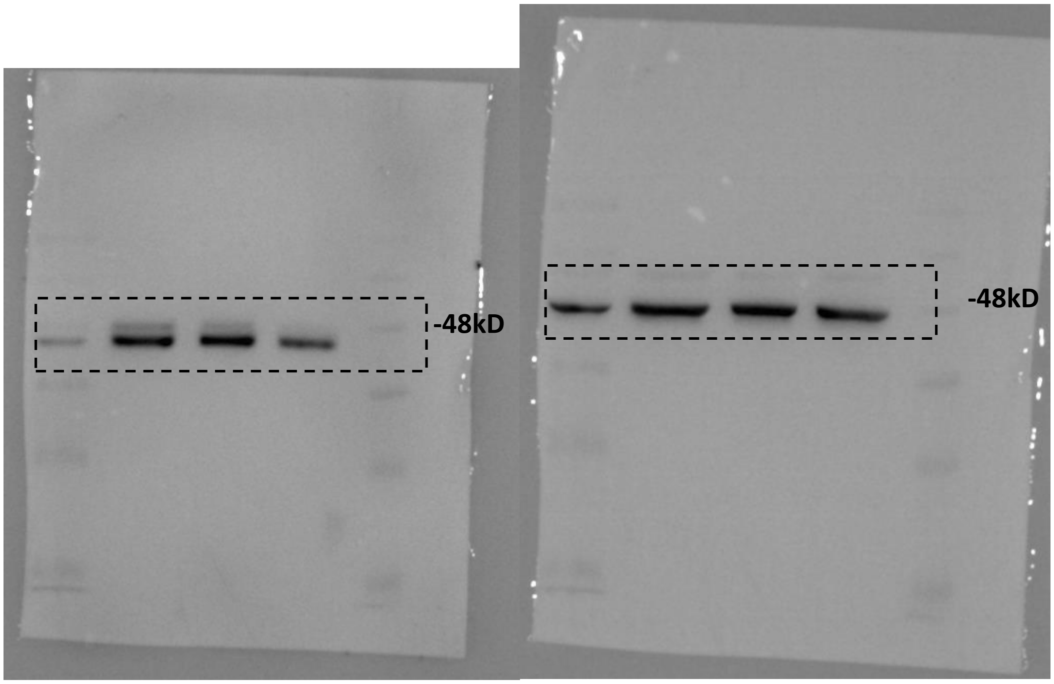

Supplementary Figure 5

**-pHSP27**

35kD-

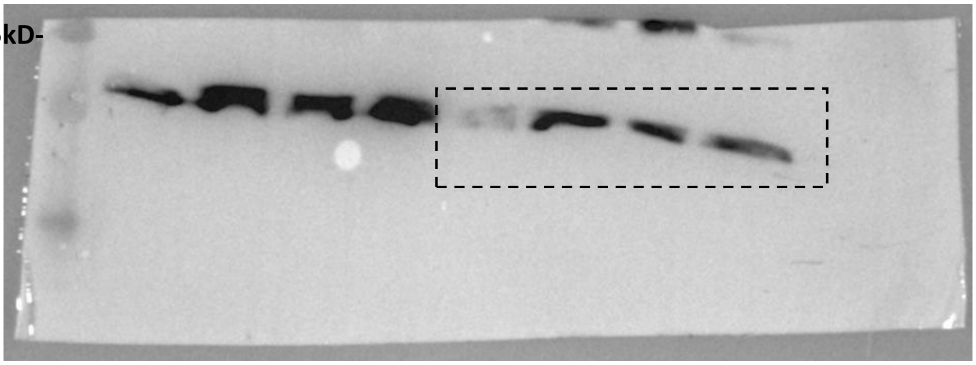

**-c-Jun**

48kD-

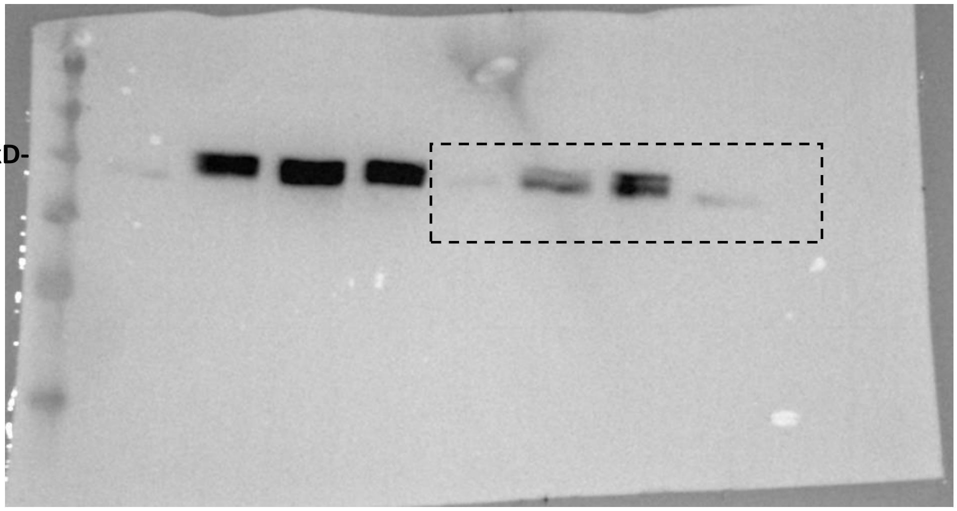

**-b-tubulin**

63kD-

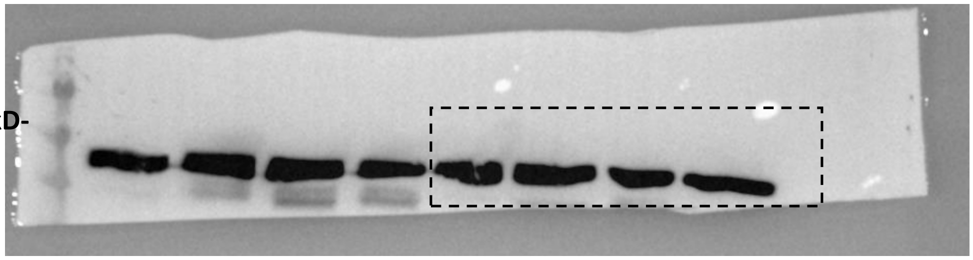

**Supplementary Figure 6**

**-pJNK**

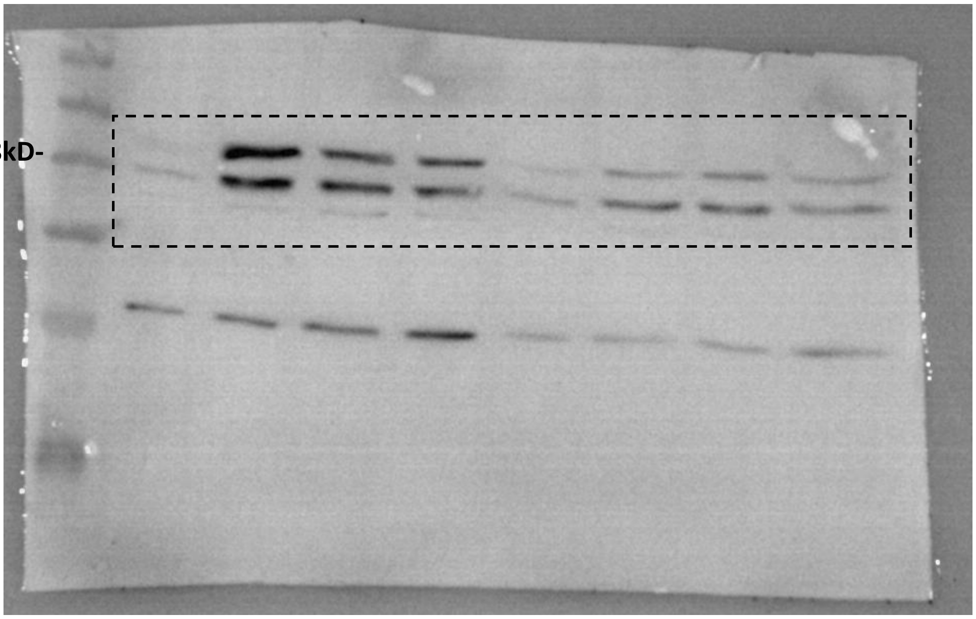


**-p-p38 MAPK**

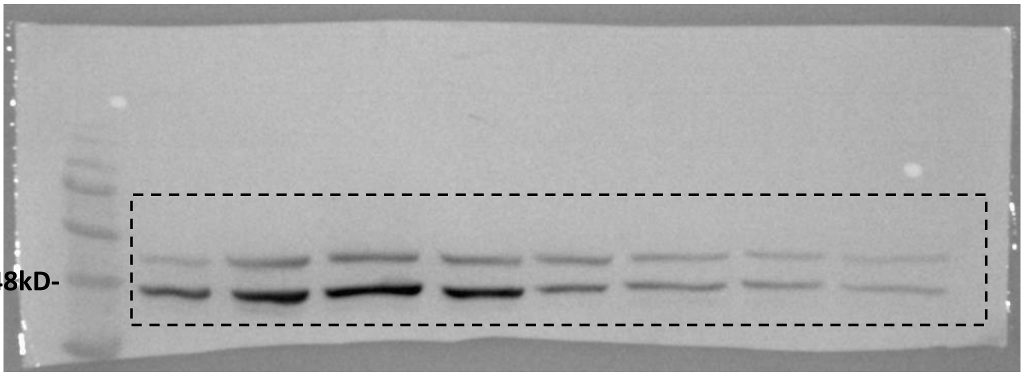


**-CDC42 TOTAL**

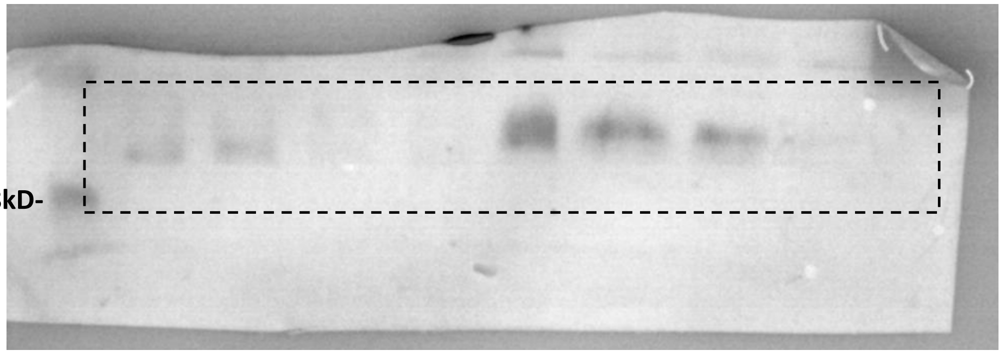


**-b-actin**

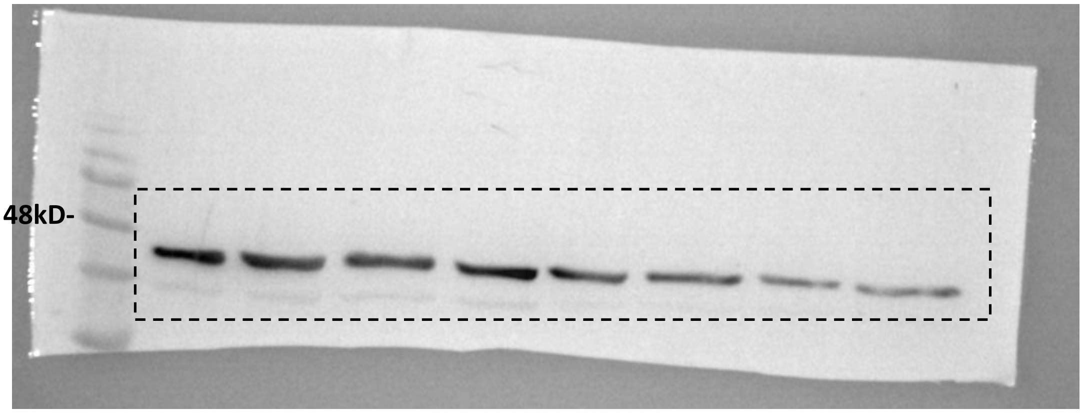


**Supplementary Table 1**

**Supplementary Table 1. Primers used for qPCR**

| Primer Name | Primer sequence |
|---|---|
| ***β-actin*** | Forward: 5'-CGAGCACAGAGCCTCGCCTTTGCC-3'<br>Reverse: 5'-TGTCGACGACGAGCGCGGCGATAT-3' |
| ***Slug*** | Forward: 5'-TGGTTGCTTCAAGGACACAT-3'<br>Reverse: 5'- GCAGATGAGCCCTCAGATTT-3' |
| ***Snail*** | Forward: 5'-AATCGGAAGCCTAACTACAAG-3'<br>Reverse: 5'-AGGAAGAGACTGAAGTAGAG-3' |
| ***Twist*** | Forward: 5'-CCTCTACCAGGTCCTCCAGA-3'<br>Reverse: 5'-ATCCTCCAGACCGAGAAGG-3' |
| ***Rac-1*** | Forward: 5'-AACCAATGCATTTCCTGGAG-3'<br>Reverse: 5'-CAGATTCACCGGTTTTCCAT-3 |
| ***cdc42*** | Forward: 5'-GCCCGTGACCTGAAGGCTGTCA-3'<br>Reverse: 5'-TGCTTTTAGTATGATGCCGACACCA-3 |

### SUPPLEMENTARY MATERIALS AND METHODS

#### Alamar Blue Assay

Cancer cells were subjected to a cell viability test prior- and post- compression using AlamarBlue reagent (Thermo), according to the manufacturer's instructions. %Difference in AlamarBlue Reagent was calculated in triplicates for each condition.
